# FH variant pathogenicity promotes purine salvage pathway dependence in kidney cancer

**DOI:** 10.1101/2022.08.15.504023

**Authors:** Blake R. Wilde, Nishma Chakraborty, Nedas Matulionis, Stephanie Hernandez, Daiki Ueno, Michayla E. Gee, Edward D. Esplin, Karen Ouyang, Keith Nykamp, Brian Shuch, Heather R. Christofk

## Abstract

The tricarboxylic citric acid cycle enzyme fumarate hydratase (FH) is a tumor suppressor. When lost in cells, its substrate fumarate accumulates to mM levels and drives oncogenic signaling and transformation. Germline alterations lead to an autosomal dominant condition known as hereditary leiomyomatosis and renal cell cancer (HLRCC) where patients are predisposed to various benign tumors and an aggressive form of kidney cancer. *FH* alterations of unclear significance are frequently observed with germline testing; thus, there is an unmet need to classify *FH* variants by their cancer-associated risk, allowing for screening, early diagnosis and treatment. Here we quantify catalytic efficiency of 74 FH variants of uncertain significance. Over half were enzymatically inactive which is strong evidence of pathogenicity. We generated a panel of HLRCC cell lines expressing FH variants with a range of catalytic activities, then correlated fumarate levels with metabolic features. We found that fumarate accumulation blocks purine biosynthesis, rendering FH-deficient cells reliant on purine salvage to maintain purine nucleotide pools. Genetic or pharmacologic inhibition of the purine salvage pathway reduced HLRCC tumor growth *in vivo*. Together, these findings suggest pathogenicity of many patient-associated *FH* variants and reveal purine salvage as a targetable vulnerability in FH-deficient tumors.

**Statement of Significance:** This study functionally characterizes patient-associated FH variants with unknown significance for pathogenicity. This study also reveals nucleotide salvage pathways as a targetable feature of FH-deficient cancers, which are shown to be sensitive to the purine salvage pathway inhibitor 6-mercaptopurine. This presents a new rapidly translatable treatment strategy for FH-deficient cancers.

## Main

Fumarate hydratase (FH) catalyzes the reversible hydration of fumarate to L-malate in the tricarboxylic citric acid cycle (TCA). Germline inactivating mutations in *FH* are associated with hereditary leiomyomatosis and renal cell cancer (HLRCC), a condition that predisposes patients to benign uterine and cutaneous leiomyomas and a unique form of renal cell carcinoma (RCC) with early onset and aggressive behavior prompting lifelong screening^1–3^. Hereditary *FH*-mutant cancers conform to Knudson’s “two-hit” model for tumorigenesis, where loss of heterozygosity (LOH) provides a “hit”—that is, loss of the wildtype allele in combination with a germline inactivating mutation are initiating steps in oncogenic transformation^4,5^. Consistent with this idea, biallelic somatic mutations in *FH* have also been observed in histologically similar forms of RCC^6,7^.

Sporadic or germline inactivating mutations in *FH* result in the accumulation of fumarate, a *bona fide* oncometabolite^8^. Though the mechanisms that underlie fumarate-driven transformation are still unknown, a combination of factors are likely necessary including: 1) inhibition of α-ketoglutarate-dependent dioxygenases resulting in global changes in DNA and histone methylation and stabilization of hypoxia-inducible factors^9–11^, 2) increased anabolic metabolism to support tumor growth^12^, and 3) dysregulation of signaling pathways that counter metabolic stress and drive proliferation^8^. Fumarate accumulation is also associated with broad metabolic rewiring, yet it remains unclear which pathway alterations are necessary to support tumor growth or are potential liabilities for cell stress/death in FH-deficient cancers.

*FH* alterations are more common than previously thought, with 0.6% of patients having a variant of unknown significance on germline testing and a possible increased prevalence of kidney cancer^13^. This highlights the need to properly classify variants according to their risk for kidney cancer and follow patients appropriately. Here we characterized 74 *FH* variants of uncertain significance (VUS) including some with conflicting interpretations of pathogenicity (CI), collectively VUS/CI. We quantified catalytic efficiencies and assessed multimerization status of each variant. Nearly half were enzymatically inactive, which is strong evidence of pathogenicity. These results help to elucidate variant pathogenicity and serve as a catalogue for interrogating fumarate-dependent changes in cells. To gain insight into the metabolic consequences of fumarate accumulation, we investigated correlations between fumarate accumulation and altered metabolism in HLRCC cells. We found that FH-deficient cells do not synthesize purines through *de novo* IMP biosynthesis but instead rely on purine salvage for nucleotide biosynthesis, thus suggesting a potential targetable metabolic vulnerability in HLRCC-associated kidney cancer.

## Results

### Patient presentation of FH variant of uncertain significance p.Met151Lys suggests pathogenicity

ClinVar reports information about genomic variation and pathogenicity by aggregating information from clinical testing laboratories. ClinVar catalogues *FH* genomic variants as associated with HLRCC and/or fumarase deficiency, which is the biallelic inheritance of loss-of-function mutations in FH. Approximately half of all *FH* variants listed in ClinVar lack a clear interpretation of pathogenicity, and thus are VUS/CI (**Fig. 1A** and Supplementary Fig. S1A). While 71% of all HLRCC-associated *FH* variants are missense mutations, only 22% of those have been interpreted as benign or pathogenic (Supplementary Fig. S1B), leaving many HLRCC-associated *FH* missense variants classified as VUS/CI.

**Fig. 1:**
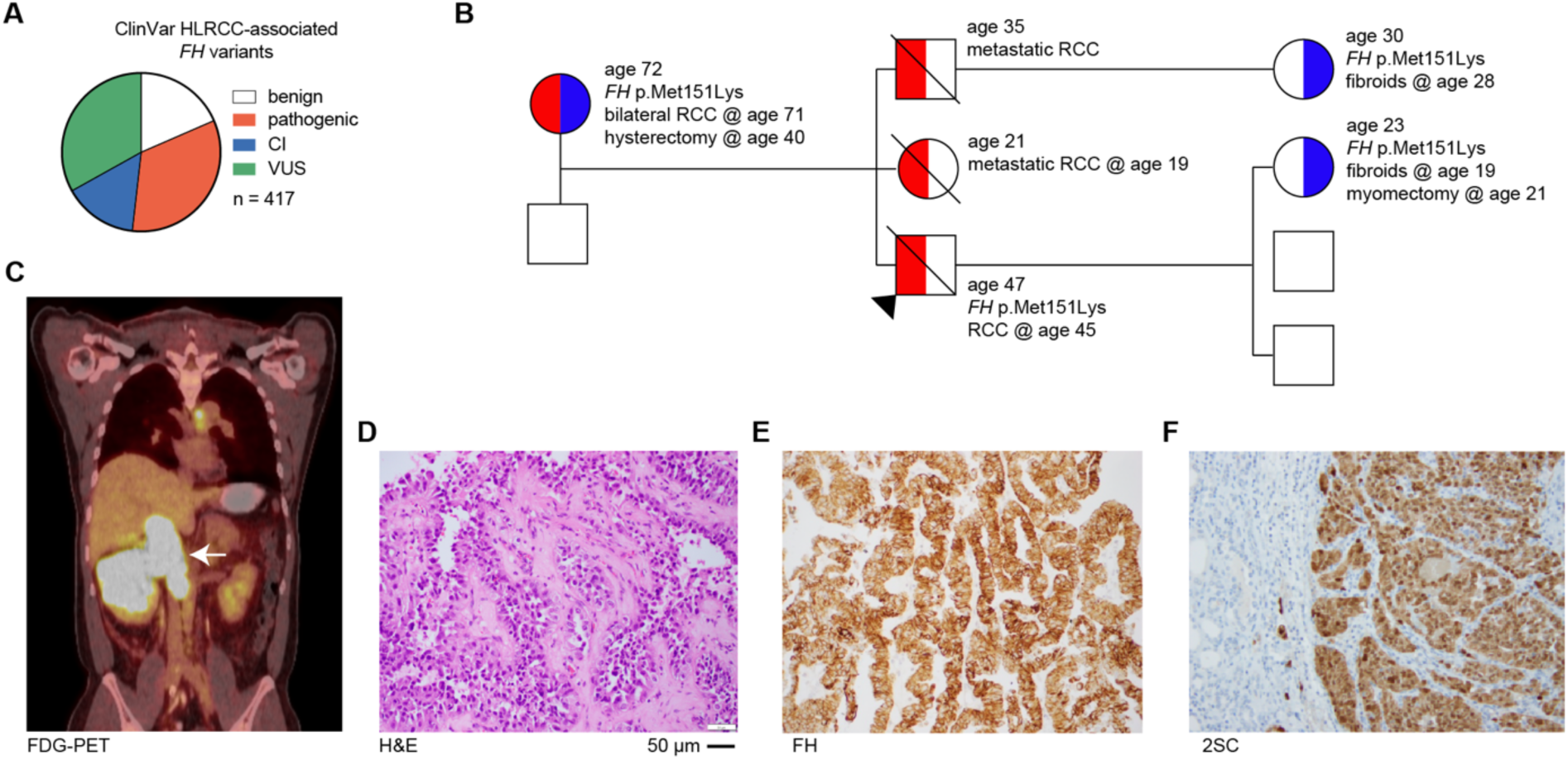
Patient presentation with germline *FH* variant of uncertain significance, p.Met151Lys, suggests pathogenicity. (**A**) Pie chart showing the proportion of HLRCC-associated *FH* variant classification as either benign (or likely-benign), pathogenic (or likely-pathogenic), conflicting interpretation of pathogenicity (CI), or variant of uncertain significance (VUS) in June 2022. (**B**) Family history of RCC patient is suspicious of HLRCC. Proband is indicated by black arrow. Colors indicate HLRCC-associated morbidities – RCC (red) and fibroids/hysterectomy/myomectomy (blue). Age is listed for each individual as well as ages at which morbidities manifested or procedures conducted. The presence of *FH* p.Met151Lys is also indicated. The probands siblings were not tested for *FH* alterations. (**C**) [^18^F]-fluorodeoxyglucose PET/CT scan of patient indicating right renal mass as indicated by white arrow. (**D**) H&E staining of renal mass demonstrates a typical HLRCC-associated kidney cancer morphology, including eosinophilic papillary architecture with classical peri-nucleolar halos and prominent organiophilic nucleoli. (**E**) Immunohistochemistry demonstrates the renal mass is FH positive. (**F**) 2-succinylcysteine (2-SC) is diffusely positive in the cytoplasm and nuclei.

Patients with *FH* variants classified as VUS/CI need clarity on whether they have HLRCC, especially so they can begin screening for RCC during adolescence. For example, a 47-year old male with a family history of RCC who was not previously screened presented symptomatically with a large renal mass (**Fig. 1B**). PET/CT scanning with ^18^F-fluorodeoxyglucose showed a large right renal mass with tumor thrombus in the inferior vena cava extending just below the edge of the diaphragm (**Fig. 1C**). Histology showed suspicious HLRCC features including eosinophilic papillary architecture with classic peri-nucleolar halos and prominent orangiophilic nucleoli (**Fig. 1D**). Immunohistochemistry showed retention of FH protein expression (**Fig. 1E**); however, 2-succinyl-cysteine (2SC) staining, a biomarker of FH-deficient tumors, showed diffusely positive cytoplasmic and nuclear staining (**Fig. 1F**). These results are consistent with a loss-of-function mutation in *FH*. Genetic testing showed a heterozygous germline alteration in *FH*, p.Met151Lys. The proband’s mother, one child, and a niece are also carriers of *FH* p.Met151Lys (M151K) and have HLRCC-associated morbidities including fibroids and RCC (**Fig. 1B**). There are conflicting interpretations of pathogenicity for this variant, arguing that despite the strong clinical evidence of pathogenicity, this polymorphism should not inform clinical decision making. This case illustrates the urgent need to interpret *FH* VUS/CI for pathogenicity so family members are assessed for *FH* variant status, and when indicated, begin regular screening for kidney cancer.

### A catalogue of 74 HLRCC-associated FH variants

We used biochemical analyses to classify FH-M151K and other *FH* VUS/CI. We expressed and purified recombinant FH from *E. coli* (**Fig. 2A**) and developed a spectroscopic assay for quantifying FH catalytic activity in the forward (hydration of fumarate) and reverse (dehydration of malate) directions. As expected, several variants reported as pathogenic in ClinVar had no quantifiable enzymatic activity (**Fig. 2B**). We extended this analysis to five VUS/CI. Three variants, pSer187Leu (S187L), p.Arg350Trp (R350W), and p.Glu376Pro (Q376P), were inactive in both directions (**Fig. 2C**). p.Pro174Arg (P174R) had a higher K_m_ for fumarate and lower k_cat_ and catalytic efficiencies for both fumarate and malate (**Fig. 2C** and Supplementary Fig. S2A-C). p.Ala70Thr (A70T) had higher K_m_ and k_cat_ for fumarate; however, catalytic efficiencies for both fumarate and malate were no different than wildtype enzyme (**Fig. 2C** and Supplementary Fig. S2D-F). These studies reveal strong evidence of pathogenicity for the inactive VUS/CI: S187L, R350W, and Q376P^14^.

**Fig. 2:**
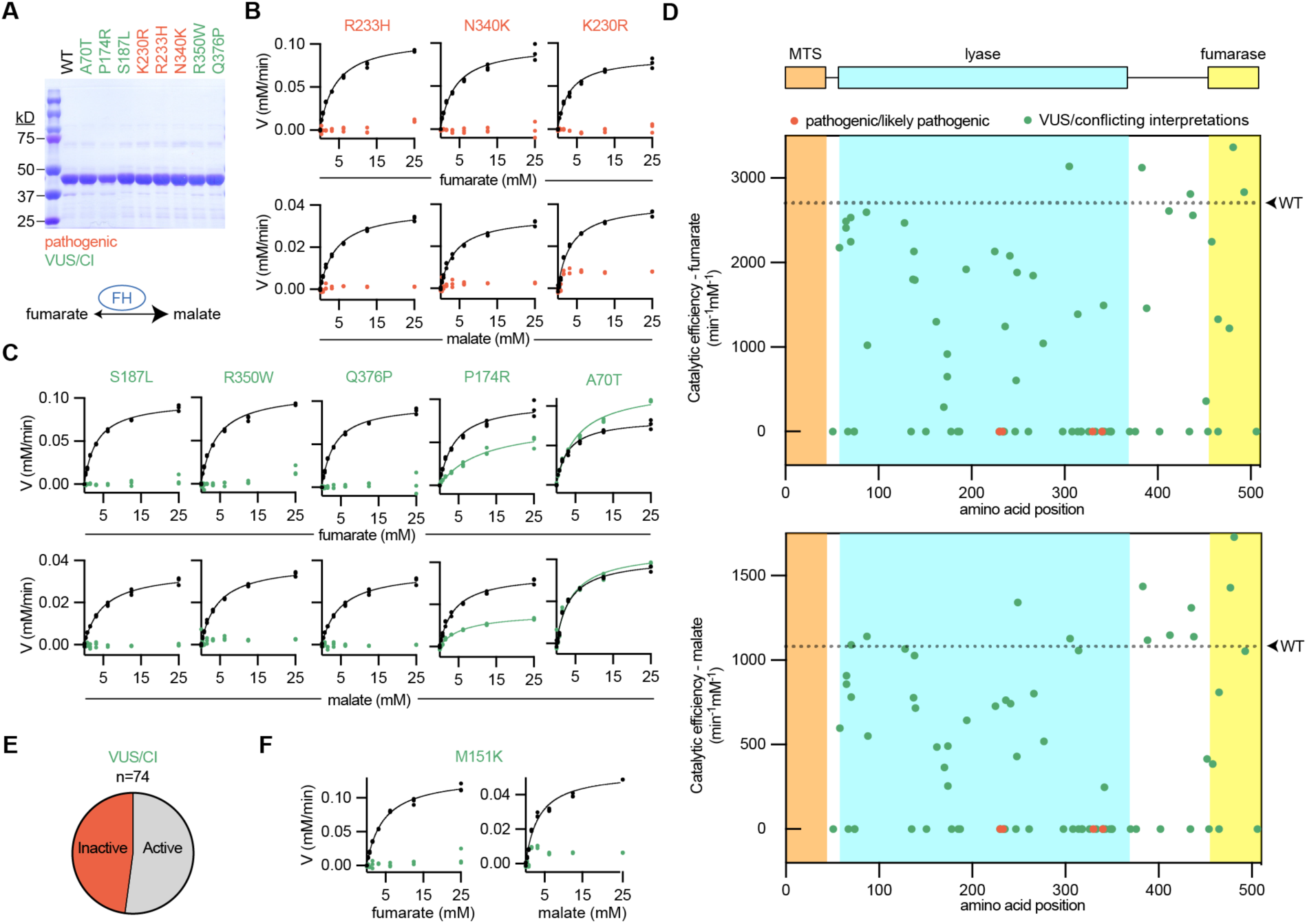
Biochemical classification of 74 patient *FH* variants provides strong evidence of pathogenicity for many VUS/CI. (**A**) Recombinant FH was produced and purified from *E. coli* as shown by SDS-PAGE gels stained by Coomassie. Each variant is colored according to the associated interpretation of pathogenicity: pathogenic (red) and VUS/CI (green). For (**B**-**D**), recombinant FH was incubated with increasing concentrations of substrate, fumarate (top) or malate (bottom), and the reaction progress was quantified. Wildtype (WT) FH was run simultaneously for each variant, as indicated by the black symbols/lines. (**B**) All pathogenic variants could not be modeled by Michaelis-Menten kinetics, indicating that these variants are inactive. Michaelis-Menten plots from wildtype enzyme are indicated by black symbols/lines. (**C**) VUS/CI have a range of enzymatic activities. (**D**) FH with annotated domains and graphs indicating the catalytic efficiencies of 74 FH variants for fumarate (top) and malate (bottom). For each graph, WT catalytic efficiency is indicated as a dotted line. No variants in the mitochondrial targeting sequence (MTS) were tested because the recombinant FH lacked the first 44 amino acids to improve solubility and purification. (**E**) Pie chart showing the proportion of VUS/CI that were inactive or active (including partial activity). (**F**) Graphs showing that M151K (product of *FH* p.Met151Lys described in Fig. 1) is enzymatically inactive.

Additional *FH* variants were included in our analyses if they met the following criteria: (1) observed in at least one patient ordering the FH test; (2) resulted in a missense change; and (3) had a Sherloc point score > 1.5 pathogenic points^15^. Thus, we extended our biochemical characterization to a prioritized list of 74 FH VUS/CI (**Fig. 2D** and Supplementary Fig. S3-S5), which represents approximately half of all VUS/CI missense variants currently listed with interpretations in ClinVar. While k_cat_ for wildtype FH was variable across production lots, catalytic efficiencies were stable and made up <10% of the total variation (Supplementary Fig. S6A-C). Thus, we quantified catalytic efficiencies for all 74 VUS/CI. Catalytic efficiencies for the forward and reverse reactions correlated well (Supplementary Fig. S6D-F). Importantly, nearly half of all VUS/CI had no measurable enzymatic activity, which is strong evidence of pathogenicity for these variants (**Fig. 2E** and Supplementary Table S1). Finally, M151K, the variant carried by the proband in Fig. 1, had no measurable enzymatic activity (**Fig. 2F**). Consistent with the clinical data, our biochemical analysis argues that *FH* p.Met151Lys is a pathogenic variant.

In addition to characterizing *FH* missense variants, we also investigated *FH* p.Lys477dup (K477dup), the most common *FH* variant with frequencies as high as 1 in 195 in some populations^16^. Though compound heterozygosity involving K477dup may be pathogenic in the context of autosomal recessive fumarase deficiency^17,18^, the presence of K477dup alone does not correlate with increased risk for RCC^13,19^. We therefore reasoned that K477dup enzymatic activity levels might be used as a potential benchmark for evaluating pathogenicity of FH variants. We thus purified K477dup and measured its catalytic activity relative to wildtype enzyme. It has diminished catalytic efficiency for fumarate and near wildtype catalytic efficiency for malate (Supplementary Fig. S3-S5, S6G). Given the lack of a role in for K477dup in HLRCC, we expect that FH variants with activity greater than that of K477dup are likely benign (Supplementary Fig. S6H).

FH forms a homotetramer with four substrate-binding sites (Supplementary Fig. S7A). It is reasonable that FH alterations that disrupt multimerization would also impact enzymatic function, yet we questioned whether multimerization status is a good predictor of FH function and thus pathogenicity. We therefore visualized abundance of multimerization species using native-PAGE with Coomassie staining and quantified bands by densitometry (**Fig. 3A**). For wildtype FH, 81% is tetrameric, with little variability across gels and production lots (**Fig. 3B** and Supplementary Fig. S7B). The pathogenic variants K230R, R233H, and N340K all had disrupted multimerization, consistent with crystal structures that place the mutated residues at multimerization interfaces (**Fig. 3A** and Supplementary Fig. S7C). However, the multimerization status of some of the VUS/CI tested was more variable; for example, Q376P and R350W are both enzymatically inactive, but only Q376P has markedly disrupted multimerization (**Fig. 3B**).

**Fig. 3:**
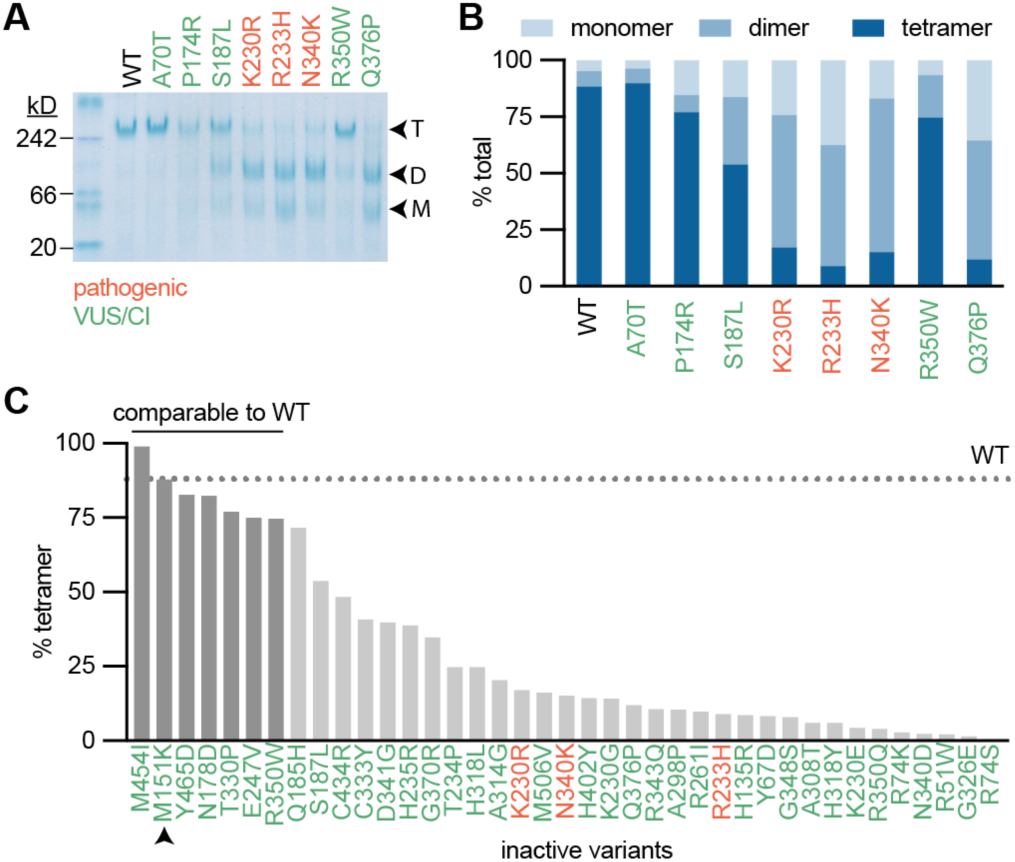
Tetramerization is necessary but not sufficient for transformation. (**A**) Recombinant FH was analyzed by Native-PAGE with Coomassie staining. Enzyme separated out to tetramers, dimers, and monomers, indicated as T, D, and M respectively. (**B**) Densitometry was used to quantify the percentage of each multimerization species for each variant. (**C**) Percent tetramerization for all variants that did not fit Michaelis-Menten kinetics in Fig. 2. Wildtype (WT) percent tetramer is indicated as a dotted line. All variants that are within 1 standard deviation of WT percent tetramer are indicated by darker gray bars.

We next analyzed all of the FH variants tested in Fig. 2 (**Fig. 3B** and Supplementary Fig. S7D-E). We observed disrupted multimerization for all variants that occurred at multimerization interfaces, including non-bonding, hydrogen-bonding, and salt bridge-forming residues. (Supplementary Fig. S8B). While there was no apparent relationship between monomeric or dimeric species and catalytic activity, there was a slight positive correlation between tetramerization and catalytic efficiency (Supplementary Fig. S8C-G), consistent with the notion that tetramerization is necessary for enzymatic activity.

A survey of all inactive variants from Fig. 2 showed significant variability in multimerization status, including M151K which had a tetramer abundance almost identical to wildtype (**Fig. 3C**). We tested whether % tetramerization could accurately classify active/inactive variants, but found that it only has modest diagnostic ability (Supplementary Fig. S8H). Together, these data suggest that FH tetramerization is necessary for enzymatic activity; however, multimerization status alone cannot predict pathogenicity.

### Defining fumarate-driven metabolism

FH inactivation is accompanied by a rise in fumarate levels and leads to broad metabolic rewiring that has been well-described by several labs (reviewed in ^8,12^). FH loss and fumarate accumulation drives the succination of L-arginine to produce argininosuccinate via reversal of the urea cycle enzyme argininosuccinate lyase^20,21^. Fumarate reacts non-enzymatically with the side chain of free or protein-incorporated cysteines to produce 2-succinyl-cysteine (2SC)^22–26^. Additionally, FH loss is associated with oxidative stress caused by the non-enzymatic succination of glutathione (GSH) to produce succinicGSH^27,28^.

We used our catalogue of FH variants to decipher how varying levels of fumarate accumulation impacts HLRCC-associated RCC cell metabolism. We generated a panel of HLRCC cells with varying levels of FH activity by expressing V5-tagged FH variants with a range of catalytic efficiencies in the HLRCC cell line NCCFH1, then we measured proliferation rates and analyzed metabolites by liquid-chromatography-mass spectrometry (LC-MS) (**Fig. 4A-B** and Supplementary Fig. S9A). Principal component analysis using the LC-MS data showed that FH variants with similar catalytic activities clustered together, demonstrating that much of the metabolic variation in this dataset is associated with FH enzymatic activity (Supplementary Fig. S9B). Fumarate/malate ratios correlated well with loss of FH catalytic efficiency and proliferation rates (Supplementary Fig. S9C-D). As expected, known fumarate-regulated metabolites, such as arginosuccinate, succinicGSH, and 2-SC, correlated well with fumarate/malate ratios (**Fig. 4C**). We next investigated how fumarate accumulation across our panel of HLRCC cells engineered to express varying levels of FH activity impacts levels of 164 metabolites routinely measured in our targeted metabolomics pipeline (**Fig. 4C**). Metabolites that significantly correlated or anti-correlated with fumarate/malate ratio with a Pearson r ≥ 0.8 or ≤ −0.8 are shown in **Fig. 4C**. Most notably, 8/10 of the metabolites that were most highly correlated with fumarate/malate ratios were involved in nucleotide biosynthesis. Additionally, metabolites in the reductive branch of the TCA cycle were negatively correlated with fumarate/malate ratios. We therefore questioned whether fumarate accumulation is sufficient to dysregulate the associated pathways.

**Fig. 4:**
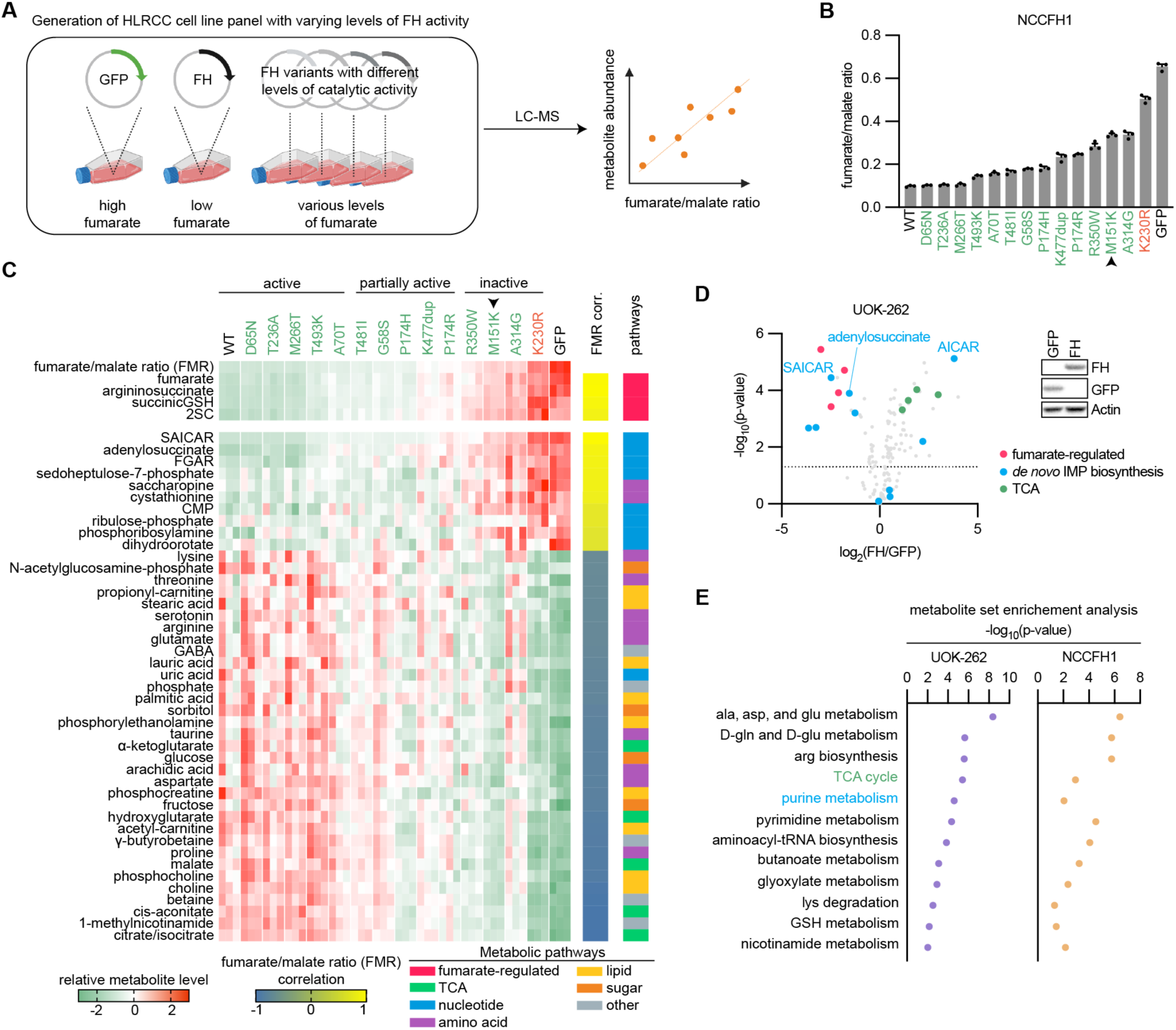
Loss of FH activity is associated with widespread metabolic changes. (**A**) Schematic depicting generation of an HLRCC cell line panel with varying levels of FH activity and fumarate. NCCFH1 cells were engineered to express GFP (control), wildtype *FH*, or *FH* variants with differing levels of catalytic activity. Metabolites were harvested and analyzed by liquid chromatography-mass spectrometry (LC-MS). Fumarate/malate ratios were calculated and correlated with other metabolites. (**B**) Fumarate/malate ratios across the panel of HLRCC cell lines engineered to have varying levels of FH activity. Pathogenic variant is indicated in red, 13 VUS/CI are indicated in green, M151K is indicated by the black arrow. **C**, Heatmap showing that fumarate/malate ratios (FMR) correlate with metabolites that have previously been linked to fumarate accumulation. This analysis was expanded to 164 targeted metabolites. Those that correlated with FMR with a Pearson r ≥ 0.8 or ≤ −0.8 are shown. (**D**) Volcano plot showing the dysregulation of metabolites in UOK-262 cells expressing FH. Fumarate-regulated metabolites, de novo IMP biosynthesis intermediates, and TCA intermediates are highlighted. Two of the most highly-altered metabolites, SAICAR and AICAR, which are sequential metabolites in *de novo* IMP biosynthesis, were regulated in opposite directions. (**E**) Metabolite set enrichment analysis was performed on HLRCC cells expressing GFP or FH. Pathways that were significant in HLRCC cell lines (p ≤ 0.05) are noted with their respective −log_10_(p-values).

### FH inactivation decreases TCA intermediates

Since FH is a TCA cycle enzyme, FH activity loss would be predicted to broadly impact TCA metabolism, even beyond the increased fumarate/malate ratio. Indeed, previous studies have found that *FH*-deficiency results in decreased TCA intermediates: α-ketoglutarate, citrate, isocitrate, and malate^29^. Consistent with this finding, our panel of cell lines with differing levels of fumarate accumulation revealed an inverse correlation between fumarate/malate ratios and TCA intermediates (**Fig. 4C**). Wildtype FH expression in another HLRCC cell line, UOK-262, led to decreased fumarate/malate ratios, a reduction in known fumarate-regulated metabolites, and increases in TCA intermediates (**Fig. 4D** and Supplementary Fig. S9E). HLRCC cells use glutamine to fuel TCA metabolism through reductive carboxylation^30^ (Supplementary Fig. S10A), which is likely a consequence of fumarate-driven PDK activity and PDH inhibition by post-translational modification^31^. We used stable-isotope labeling of glutamine and glucose to determine the relative contribution of each carbon source to TCA intermediates in conditions where fumarate is high (HLRCC cells expressing GFP) versus low (HLRCC cells expressing wildtype FH). Consistent with glutamine as the primary source of carbons for the TCA cycle in cells with elevated fumarate levels, HLRCC cells expressing GFP labeled with U-^13^C-glutamine had a higher ratio of ^13^C-labeled TCA metabolites than HLRCC cells labeled with U-^13^C-glucose (Supplementary Fig. S10B-D). FH expression in two different HLRCC cell lines, UOK-262 and NCCFH1, only slightly increased the ratio of U-^13^C-glucose-labeled TCA metabolites or had no effect on TCA metabolite glucose labeling, respectively (Supplementary Fig. S10A-C). As expected, the total levels of fumarate and succinate were decreased and malate levels were increased in HLRCC cells expressing FH (Supplementary Fig. S10A-D). While FH expression led to increased levels of intermediates within the reductive branch of the TCA cycle, no consistent changes were observed in acetyl-CoA levels.

In cells labeled with U-^13^C-glutamine, nearly all of the citrate/isocitrate, cis-aconitase, and α-ketoglutarate were labeled as m+5, consistent with glutamate metabolism through reductive carboxylation (Supplementary Fig. S10D). While cells expressing FH increased flux of carbons through oxidative decarboxylation, as evidenced by m+4 malate, we observed persistent m+5 labeling of citrate/isocitrate, cis-aconitase, and α-ketoglutarate, indicating that reductive carboxylation is the primary route of glutamine metabolism even when fumarate is low. These findings show that transient changes in fumarate levels are insufficient to alter the flow of carbons through reductive carboxylation in HLRCC cells.

### FH inactivation leads to glutathione biosynthesis

Among the metabolites that most strongly correlated with fumarate/malate ratios was cystathionine, a precursor of cysteine synthesis (**Fig. 4C**). The acquisition of cysteine is generally considered to be the rate-limiting step in glutathione biosynthesis^32,33^; thus, we hypothesized that fumarate accumulation regulates *de novo* glutathione biosynthesis. Consistent with this idea, metabolite set enrichment analysis (MSEA) revealed significant regulation of glutathione metabolism in both HLRCC cell lines expressing FH (**Fig. 4E**). We therefore analyzed metabolites involved in glutathione biosynthesis and found that FH expression in HLRCC cells decreased cystathionine and cysteine levels (Supplementary Fig. S11A), though GSH levels were unchanged. FH expression in HLRCC cells had no effect on the GSH/GSSG ratios, indicating that despite fumarate accumulation, cells with and without FH expression achieve similar levels of redox homeostasis (Supplementary Fig. S11B). Glutathione is a tripeptide consisting of cysteine, glutamate, and glycine. The reactive sulfur on the side chain of cysteine can be oxidized and broadly controls the redox status of cells. We therefore analyzed cysteine incorporation to glutathione by culturing cells with 3,3,3’,3’-^2^H-cystine, a precursor to cysteine followed by LC-MS-based metabolite analysis (Supplementary Fig. S11C). GSH and oxidized glutathione (GSSG) were labeled by cysteine, though the labeling was decreased in HLRCC cells expressing FH (Supplementary Fig. S11D). There was a slight increase in the labeling of cysteine and γ-glutamylcysteine (γ-GluCys) in UOK-262 cells expressing FH, but no change in NCCFH1 cells expressing FH. We next analyzed incorporation of glutamate in GSH by labeling cells with U-^13^C-glutamine. Consistent with the cystine-tracing experiments, GSH and GSSG were labeled, but to a lesser extent in HLRCC cells expressing FH (Supplementary Fig. S11E). FH expression slightly decreased the labeling of glutamate, but had no effect on the labeling of γ-GluCys. The lack of change in glutamine labeling of γ-GluCys together with the decreased labeling of GSH and GSSG in HLRCC cells expressing FH suggest that fumarate accumulation impacts reactions downstream of γ-GluCys to increase glutathione biosynthesis. Finally, fumarate accumulation is negatively correlated with ophthalmate, a biomarker of glutathione depletion (Supplementary Fig. S11E)^34^. Together these data suggest that the increased glutathione biosynthesis in HLRCC cells, which lack functional FH, may counter oxidative stress caused by GSH depletion through the formation of succinicGSH (Supplementary Fig. S11F).

### FH inactivation restricts de novo purine biosynthesis

Interestingly, the two metabolites that most highly correlated with fumarate/malate ratios across our panel of NCCFH1 HLRCC cells engineered to express FH variants with differing levels of enzymatic activity (**Fig. 4A**) were 1-(phosphoribosyl)imidazolecarboxamide (SAICAR) and adenylosuccinate (**Fig. 4C**), which are both substrates of adenylosuccinate lyase (ADSL)-catalyzed reactions. Consistently, wildtype FH expression UOK-262 cells led to decreased fumarate/malate ratios, a reduction in known fumarate-regulated metabolites, and decreased levels of both SAICAR and adenylosuccinate (**Fig. 4D** and Supplementary Fig. S9E). Along with SAICAR, we observed a decrease in other nucleotide biosynthesis intermediates in UOK-262 cells with wildtype FH expression. Additionally, MSEA indicated that purine metabolism is among the most highly altered pathways in HLRCC cell lines engineered to express GFP versus wildtype FH (**Fig. 4E**). Finally, analysis of co-dependency scores by the DepMap project revealed a correlation between *FH* essentiality and many genes involved in nucleotide biosynthesis, which suggests a potential functional overlap in the pathways (Supplementary Fig. S12)^35^. Together, these data indicate that FH catalytic activity may regulate nucleotide biosynthesis.

Surprisingly, we found that AICAR, a product of SAICAR metabolism by ADSL, was the most significantly enriched metabolite in UOK-262 cells expressing FH (**Fig. 4D**). This suggests a potential uncoupling of *de novo* IMP biosynthesis in conditions where fumarate is accumulated. Fumarate is produced in purine biosynthesis by ADSL which catalyzes two reactions: 1) desuccination of SAICAR to produce AICAR and fumarate, and 2) desuccination of adenylosuccinate to produce AMP and fumarate (**Fig. 5A** and Supplementary Fig. S13A). The observation that adenylosuccinate accumulates in Fh1-knockout mouse tissues has led others to theorize that ADSL is reversed when fumarate accumulates; however, this hypothesis remains to be tested^12,36^. Thus, we used stable-isotope tracing to investigate the directionality of ADSL-catalyzed reactions. We labeled HLRCC cell lines with U-^13^C-glutamine and analyzed metabolites by LC-MS. Glutamine provides carbons to both fumarate and aspartate with predominant isotopomers m+4 and m+3 respectively (**Fig. 5A-B** and Supplementary Fig. S13B). While neither AIR nor AICAR showed any mass shifts consistent with glutamine-derived carbons, we detected m+4, and to a lesser extent m+3, labeling of SAICAR, suggesting that fumarate contributes to synthesis of SAICAR (**Fig. 5B** and Supplementary Fig. S13B). Likewise, neither IMP nor AMP were labeled, yet we detected m+4 labeling of adenylosuccinate, consistent with fumarate reacting with AMP to produce adenylosuccinate (Supplementary Fig. S13C). These data indicate that fumarate accumulation drives the succination of AICAR and AMP via the reversal of ADSL, which likely blocks the flow of carbons through *de novo* IMP biosynthesis and AMP synthesis.

**Fig. 5:**
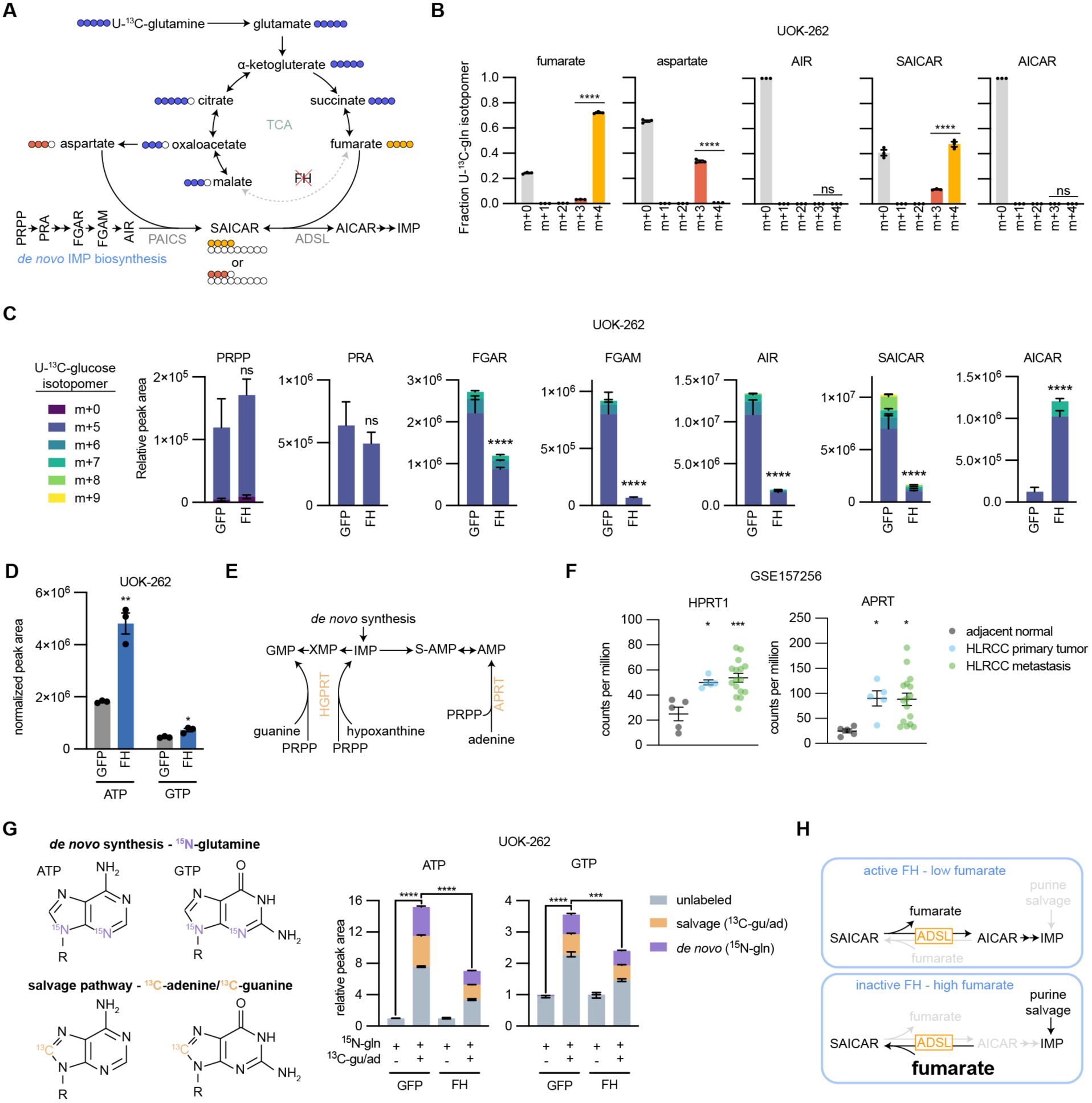
Loss of FH activity disrupts *de novo* purine biosynthesis. (**A**) Schematic representing the flow of carbons in the TCA cycle and *de novo* IMP biosynthesis from U-^13^C-glutamine in HLRCC cells. Fraction of ^13^C-labeled isotopomers for (**B**) fumarate, aspartate, AIR, SAICAR, and AICAR. (**C**) Relative peak areas for *de novo* IMP biosynthesis intermediates in UOK-262 cells treated with U-^13^C-glucose. Bars are colored by the relative amount of individual ^13^C isotopomers. (**D**) Normalized peak areas for ATP and GTP. (**E**) Schematic of the purine salvage pathway. (**F**) Expression of APRT and HPRT1 in HLRCC patient primary tumors, metastases, and adjacent normal kidney tissue. Data is publicly available at NCBI Gene Expression Omnibus GSE157256. (**G**) Depiction of how ATP and GTP are labeled by amide-^15^N-glutamine, 8-^13^C-adenine, and 8-^13^C-guanine. UOK-262 cells were treated for 6 hours with 4 mM amide-^15^N-glutamine, 50 µM 8-^13^C-adenine, and 50 µM 8-^13^C-guanine, and ATP and GTP were analyzed by LC-MS for mass shifts corresponding to *de novo* synthesis (purple) or salvage pathway (orange). (**H**) Summary of how FH loss impacts purine biosynthesis. In cells without active FH, fumarate accumulates which drives the succination of AICAR by ADSL to produce SAICAR (reverse reaction). This prevents *de novo* IMP biosynthesis, leaving the cells reliant on the salvage pathway to support purine synthesis.

We next examined how purine biosynthesis is affected by fumarate accumulation and ADSL reversal. HLRCC cells expressing GFP or FH were labeled with U-^13^C-glucose, and metabolites were analyzed by LC-MS. Phosphoribosyl pyrophosphate (PRPP) is the initiating metabolite in *de novo* IMP biosynthesis and serves as the ribose base for nucleotide synthesis; atoms from glycine, aspartate, glutamine, CO_2_, and 10-formyltetrahydrofolate are also used to construct IMP. We found that PRPP and phosphoribosylamine (PRA) were fully labeled from U-^13^C-glucose and unchanged in abundance upon expression of FH in HLRCC cells (**Fig. 5C** and Supplementary Fig. S13D). However, levels of downstream purine biosynthesis intermediates were disrupted by FH expression: phosphoribosyl-N-formylglycineamide (FGAR), phosphoribosylformylglycineamine (FGAM), 5-aminoimidazole ribotide (AIR), and SAICAR levels were decreased, and AICAR levels were significantly increased in HLRCC cells expressing FH. These data suggest a block in conversion of SAICAR to AICAR caused by fumarate accumulation in FH-deficient HLRCC cells that can be relieved by expression of wildtype FH. Consistently, FH expression increased the abundance of ATP and GTP (**Fig. 5D** and Supplementary Fig. S13E). In HLRCC cells expressing GFP we observe m+5, m+6, m+7, m+8, and m+9 isotopomers of SAICAR, but only the m+5 isotopomer of AICAR, further suggesting that SAICAR conversion to AICAR is diminished upon fumarate accumulation (**Fig. 5C** and Supplementary Fig. S13E). FH expression in HLRCC cells leads to m+7 isotopomer detection of SAICAR and AICAR (**Fig. 5C**), consistent with ADSL conversion of SAICAR to AICAR in the absence of fumarate accumulation.

We reasoned that HLRCC cells might adapt to the altered metabolism initiated by expression of exogenous FH, thus we queried whether altered purine metabolism is a direct effect of FH expression in HLRCC cells versus an effect of passage-mediated cell adaptation. We found that NCCFH1 cells infected with lentiviral FH have widescale metabolic alterations compared to control cells, but that differential levels of many metabolites are diminished over time (Supplementary Fig. S14A). As might be expected for direct effects of FH expression in HLRCC cells, exogenous FH expression led to a stable decrease in fumarate/malate ratios and decreased levels of known fumarate-regulated metabolites that remained unchanged over several passages (Supplementary Fig. S14B). Likewise, FH expression decreased SAICAR and adenylosuccinate levels in a manner that was independent of passage number, suggesting that altered purine metabolism is also a direct effect of FH expression in HLRCC cells. Importantly, this sustained effect on purine metabolism occurred despite passage-dependent changes in FH protein levels and cell proliferation rates (Supplementary Fig. S14C-E). We confirmed the sustained decrease in SAICAR levels in FH-expressing UOK-262 cells, which was also stable across passages despite changes in cell proliferation (Supplementary Fig. S14F-I).

Given our findings that FH loss-of-function causes accumulation of SAICAR and other purine intermediates (**Fig. 4C**) and that fumarate accumulation drives ADSL activity towards SAICAR production while blocking generation of AICAR (**Fig. 5A-D**), we expected that HLRCC cells with accumulated fumarate and decreased *de novo* IMP biosynthesis would be more reliant on purine salvage pathways for nucleotide biosynthesis and proliferation. Under replication stress, human cells can synthesize nucleotides through the purine salvage pathway that consists of the enzymes adenine phosphoribosyltransferase (APRT) and hypoxanthine-guanine phosphoribosyltransferase (HGPRT), which catalyze reactions between PRPP and nucleosides to generate AMP, GMP, and IMP (**Fig. 5E**). We analyzed publicly available gene expression data and found that HLRCC primary tumors and metastases express APRT and HPRT1 at higher levels than adjacent normal kidney tissue (**Fig. 5F**).

To get a sense for relative activities of APRT and HGPRT in HLRCC cells, we used a co-labeling strategy to measure contribution of *de novo* and salvage pathways to purine biosynthesis. We labeled HLRCC cells for 6 hours with amide-^15^N-glutamine and ^13^C-adenine+^13^C-guanine, then used LC-MS to decipher the labeling patterns. We found that ^13^C-adenine+^13^C-guanine dramatically increased levels of purine nucleotides in UOK-262 and NCCFH1 HLRCC cells (**Fig. 5G** and Supplementary Fig. S15A-C). ^13^C-adenine+^13^C-guanine stimulated purine labeling through both salvage and *de novo* pathways in a manner that was reduced by exogenous FH expression. Together these data demonstrate that fumarate accumulation in HLRCC cells blocks purine synthesis, rendering the cells reliant on purine bases to maintain nucleotide pools (**Fig. 5H**).

We next investigated whether pyrimidine biosynthesis is similarly disrupted by FH loss. We found that FH expression in HLRCC cells increased the levels of aspartate, carbamoyl-aspartate, and dihydroorotate; however, we did not detect changes in UMP levels or labeling (Supplementary Fig. S16A-B). Additionally, we did not detect FH-dependent changes in sensitivity to the pyrimidine salvage inhibitor, gemcitabine. Together these findings suggest that pyrimidine nucleotide synthesis is not affected by FH loss (Supplementary Fig. S16C-D), and highlight the specificity of the fumarate-driven block in purine biosynthesis.

### Targeting purine salvage reduces HLRCC tumor growth

Given that fumarate accumulation leads to a dependence on the purine salvage pathway to maintain purine nucleotides, we reasoned that inhibiting purine salvage might restrict HLRCC-associated kidney cancer growth. Thus, we used a lentiviral strategy to express Cas9 and sgRNAs targeting AAVS1 (control), APRT, and HPRT1 to disrupt expression of the enzymes (**Fig. 6A** and Supplementary Fig S17A). NCCFH1 cells displayed efficient knockout of both enzymes; however, in UOK-262 cells, the expression of HGPRT expression was restored by passaging cells, suggesting that a population of HGPRT-expressing cells out-competed knockout cells (Supplementary Fig. S17B). Despite the differences in knockout efficiencies, we observed decreased labeling of AMP, ADP, and ATP from ^13^C-adenine, and for NCCFH1 cells, we also observed decreased labeling of GMP, ADP, and ATP from ^13^C-guanine (**Fig. 6B**). When grown in human plasma-like media supplemented with adenine and guanine, sgAPRT/sgHGPRT1-exprssing cells had decreased proliferation rates (**Fig. 6C**). Further, NCCFH1 cells expressing sgAPRT/sgHPRT1 grew much slower as subcutaneous tumor xenografts in NSG mice than cells expressing sgAAVS1 (**Fig. 6D** and Supplementary Fig. S17C). These data support an important role for purine salvage in promoting HLRCC tumor growth *in vivo*. We sought to confirm these results using another HLRCC cell line, but in our hands, UOK-262 cells are inconsistent in their ability to form tumor xenografts. We therefore established a new HLRCC cell line from a patient-derived xenograft (xp-152)^37^, which we named xp-152-cl. We confirmed xp-125-cl cells lack FH activity by tracing 13C-glutamine metabolism, which showed no detectable conversion of m+4 fumarate to m+4 malate, but instead all malate was of the m+3 isotopologue, derived from reductive carboxylation (Supplementary Fig. S17D). Further, FH expression in xp-152-cl cells increased malate levels and decreased levels of fumarate and fumarate-regulated metabolites (Supplementary Fig. S17E). As with UOK-262 cells, expression of sgAPRT/sgHPRT1 in xp-152-cl cells decreased APRT but not HGPRT expression and diminished labeling of AMP, ADP, and ATP from ^13^C-adenine (**Fig. 6E-F**). Much like with the NCCFH1 cell line, expression of sgAPRT/sgHPRT1 decreased growth of xp-152-cl cell line-derived tumor xenografts (**Fig. 6G**), confirming the importance of the purine salvage pathway for HLRCC tumor growth.

**Fig 6:**
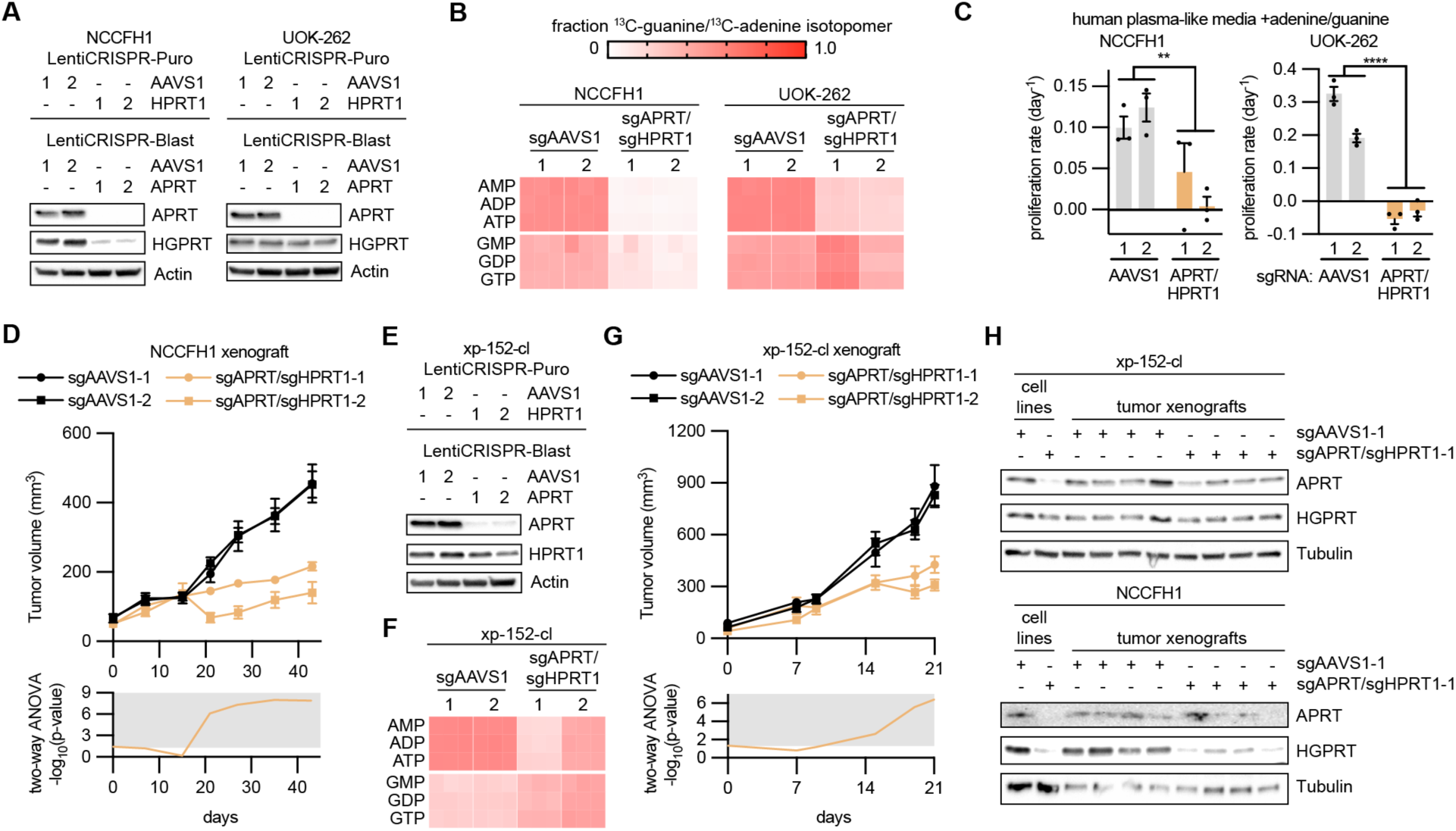
Purine salvage pathway enzymes promote HLRCC tumor growth. (**A**) Immunoblots evaluating APRT and HGPRT expression in NCCFH1 and UOK-262 cells infected with LentiCRISPR-Puro and LentiCRISPR-Blast containing sgRNAs targeting AAVS1 (control), APRT, and HPRT1. (**B**) Fraction of m+1 purine nucleotide isotopologues resulting from 6 hour treatment with 50 µM 8-^13^C-adenine and 50 µM 8-^13^C-guanine. (**C**) Proliferation rates in cells expressing sgAAVS1 or sgAPRT/sgHPRT1 at treated with human plasma-like media supplemented with 50 µM adenine and 50 µM guanine. (**D**) Volume of NCCFH1 tumor xenografts expressing sgAAVS1 or sgAPRT/sgHPRT1 as determined by caliper measurements (n = 10). Two-way ANOVA was performed for each time point and −log_10_(p-value) is plotted below, with significant values (p ≤ 0.05) falling into the gray region. (**E**) Immunoblots evaluating the expression of APRT and HGPRT in xp-152-cl cells infected with LentiCRISPR-Puro and LentiCRISPR-Blast containing sgRNAs targeting AAVS1 (control), APRT, and HPRT1. (**F**) Fraction of m+1 purine nucleotide isotopologues resulting from 6 hour treatment with 50 µM 8-^13^C-adenine and 50 µM 8-^13^C-guanine. (**G**) Volume of xp-152-cl tumor xenografts expressing sgAAVS1 or sgAPRT/sgHPRT1 as determined by caliper measurements (n = 8). Two-way ANOVA was performed for each time point and −log_10_(p-value) is plotted below, with significant values (p ≤ 0.05) falling into the gray region. (**H**) Immunoblots evaluating the expression of APRT and HGPRT in lysates from NCCFH1 and xp-152-cl cell lines and corresponding tumor xenograft lysates.

An important point is that both sets of sgAPRT/sgHPRT1-expressing tumors exhibited residual growth. Because we used a lentiviral system to knockout APRT and HPRT1, we in effect generated a heterogenous pool of knockout cells, where most cells were knockout, yet there were persisting cells that did in fact express APRT and HPRT1. Thus, we analyzed the resulting tumors for APRT and HGPRT expression and observed nearly equal levels of APRT and HPRT1 to sgAAVS1-expressing tumors, suggesting that the residual tumor growth was a result of outgrowth of wild type cells (**Fig. 6H**).

We next decided to target purine salvage pharmacologically with 6-mercaptopurine (6-MP), an analogue of hypoxanthine and potent inhibitor of APRT and HGPRT (**Fig. 7A**). We found that UOK-262 and NCCFH1 HLRCC cells were sensitive to 6-MP treatment in a FH-dependent manner: FH expression increased the EC50 of 6-MP by an order of magnitude in both HLRCC cell lines (**Fig. 7B-C**). 6-MP drives cell cycle arrest in HLRCC cells, but does not initiate apoptosis in HLRCC cells (Supplementary Fig. S18A-B). Exogenous nucleosides were unable to rescue cells from 6-MP-driven decreases in proliferation, confirming the importance of purine salvage for cell proliferation (Supplementary Fig. S18C). Finally, we found that clinically-relevant doses of 6-MP decrease the growth of NCCFH1-derived and xp-157-cl-derived tumor xenografts *in vivo* (**Fig. 7E**). Together these data demonstrate that fumarate-driven reversal of ADSL enzymatic activity towards SAICAR and adenylosuccinate, decreases *de novo* IMP biosynthesis, and renders HLRCC cells more reliant on purine salvage to support proliferation and tumor growth (**Fig. 7F**).

**Fig. 7:**
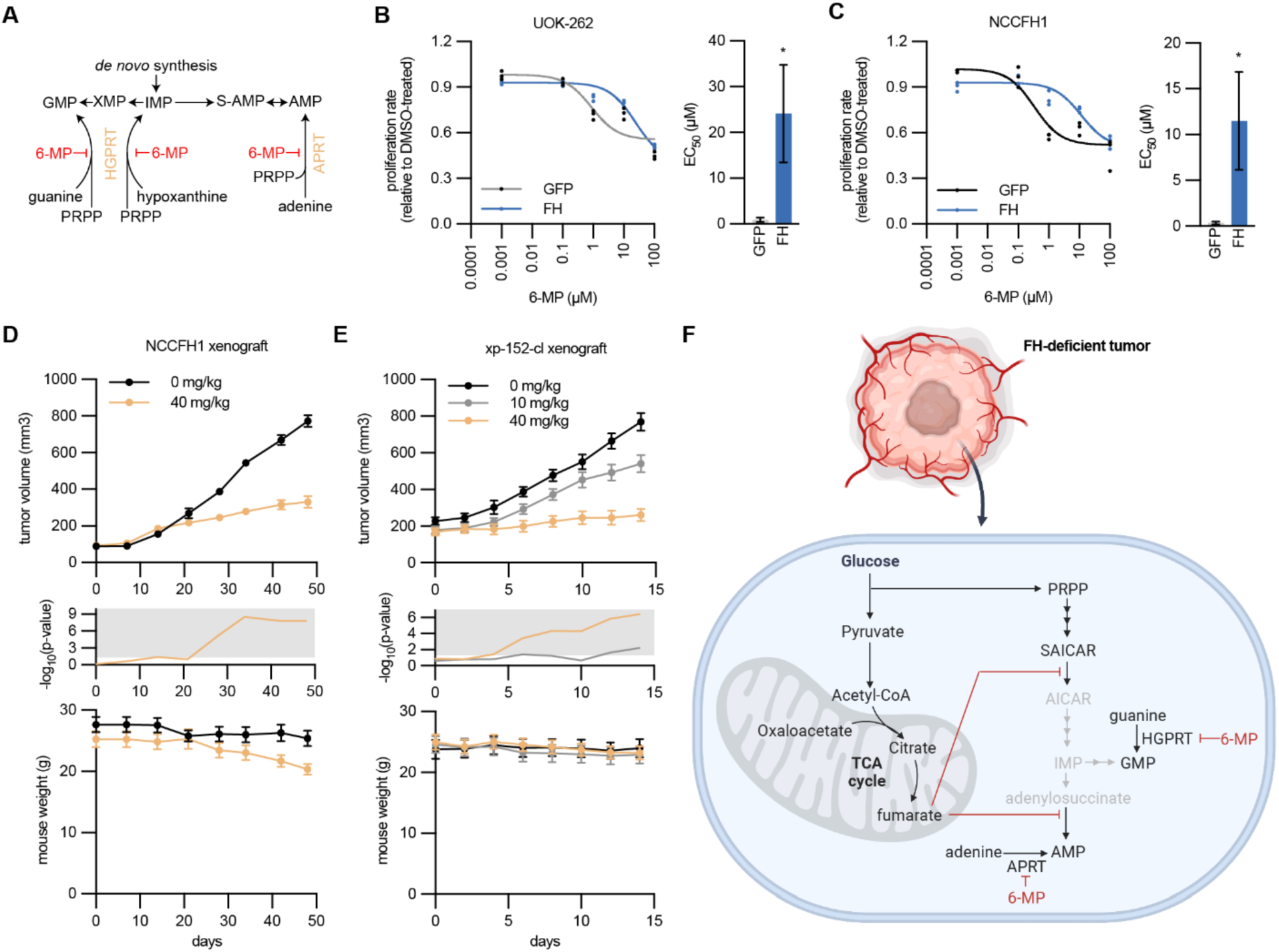
Inhibition of purine salvage by 6-mercaptopurine disrupts HLRCC tumor growth. (**A**) Schematic of the purine salvage pathway and inhibition by 6-mercaptopurine (6-MP). (**B**-**C**) Proliferation (relative to DMSO treated cells) of NCCFH1 and UOK-262 cells expressing GFP or FH and treated with varying concentrations of 6-MP. Fitted values for EC_50_ are higher for FH-expressing cells. (**D**) Volume of NCCFH1 tumor xenografts from mice treated with vehicle or 40 mg/kg 6-MP (n = 10). Student’s T-test was performed for each time point and −log_10_(p-value) is plotted in the middle, with significant values (p ≤ 0.05) falling into the gray region. Mouse weight for each time point is plotted at the bottom. (**E**) Volume of xp-152-cl tumor xenografts from mice treated with vehicle, 10 mg/kg 6-MP, or 40 mg/kg 6-MP (n = 10). Student’s T-test was performed for each time point and −log_10_(p-value) is plotted in the middle, with significant values (p ≤ 0.05) falling into the gray region. Mouse weight for each time point is plotted at the bottom. (**F**) Schematic illustrating how fumarate accumulation disrupts *de novo* purine biosynthesis, rendering FH-deficient tumor cells reliant on purine salvage for maintenance of purine nucleotides and tumor growth.

## Discussion

As genetic testing is more readily used in the clinic, there is a resultant increase in the detection of *FH* alterations, many of which have an unclear association with disease. The American College of Medical Genetics and Genomics (ACMG) recommends that VUS should not be used in clinical decision-making^14^; thus, a significant number of patients harboring these alterations may not receive regular kidney cancer screening despite suspicion that their *FH* alterations might be pathogenic. For example, in Fig. 1 we present the case of a patient with RCC. The family history is highly suspicious of HLRCC: radiological and histological analysis of HLRCC confirm FH deficiency in the tumor, and germline testing showed the presence of a *FH* variant, p.Met151Lys. The clinical data represents one strong and two supporting evidences of pathogenicity, which allows for the assertion of this variant as “likely pathogenic”. Our enzymatic assays provide an additional strong evidence of pathogenicity (PS3), and when coupled with the data in Fig. 1, supports the assertion of this variant as “pathogenic”. This case represents a scenario where the enzymatic data can be used to reinterpret the pathogenicity of a variant. Importantly, reevaluation of variants by others in the medical genetics field is necessary to change the overall interpretation of VUS/CI, and until that occurs offering screening to family members (as well as obtaining insurance authorization) would be controversial. This highlights the need to broaden our classification of patient-associated *FH* alterations in light of a potentially lethal kidney cancer syndrome where screening is cost-effective and saves lives^38^.

We often need many layers of evidence to accurately predict pathogenicity, including functional analyses. The metabolic nature of FH and its critical role in TCA metabolism made this an ideal gene to functionally interrogate. Therefore, we sought to predict the pathogenicity of *FH* variants by quantifying catalytic efficiency in cell-free assays. Of the 74 VUS/CI tested, nearly half had no measurable enzymatic activity which is strong evidence of pathogenicity, and highlights the urgency to reclassify and contact individuals with these types of *FH* VUS/CI.

Our catalogue of FH VUS/CI includes variants occurring at many different sites in the protein, including at multimerization interfaces. FH forms homotetramers with 4 active sites for fumarate/malate-binding and catalysis. Thus, we analyzed the multimerization status of all 74 VUS/CI and 3 pathogenic variants. The abundance of tetramers correlated, although weakly, with catalytic efficiencies, which is consistent with the notion that tetramerization is a prerequisite for enzymatic efficiency. However, we also observed 13 out of 74 variants where multimerization was unaltered, but catalytic efficiencies were partially or entirely compromised (Supplemental Table S1). This is highlighted by variants such as M151K (the variant encoded by the germline alteration in the patient from Fig. 1), where multimerization status is similar to wildtype, yet the enzyme is catalytically inactive. Thus, while disrupting tetramerization appears to be sufficient to restrict catalytic activity, multimerization status itself has limited value for predicting catalytic activity and associated pathogenicity alone.

Our classification of *FH* variants by enzymatic activity appears to robustly predict pathogenicity; pathogenic variants and M151K were enzymatically inactive. Our analysis of K477dup, a variant which does not confer additional RCC risk^13^, demonstrates its diminished catalytic activity (Supplementary Fig. 6G-H); this variant might be used as a potential benchmark for catalytic activity, where variants with higher activity would be sufficient to suppress tumorigenesis. However, there are some limitations to this approach. For example, these studies cannot rule out the potential that some variants, while functional in the cell-free system, may be regulated at other levels in HLRCC cells (e.g. phosphorylation). However, it is worth noting that for the *FH* variants expressed in HLRCC cell lines (Fig. 4), fumarate/malate ratios correlated with their catalytic efficiencies measured in the cell-free assays (Supplementary Fig. S9C). Another possible limitation is the lack of *in vivo* models with the variants in question that would confirm pathogenicity. We and others are working toward developing *in vivo* models of HLRCC that would better recapitulate fumarate-driven cell transformation; we expect that as these models become available, pathogenicity of many FH variants will be able to be tested *in vivo*.

Though our studies provide strong evidence of pathogenicity for inactive variants, it is more difficult to classify the variants with partial activity. For instance, is there a threshold of fumarate accumulation that is necessary for tumorigenesis? Alternatively, is there a linear relationship between fumarate accumulation and the potential for tumorigenesis? The identification of these variants with partial activity allowed us to investigate how subtle changes in fumarate accumulation might impact pro-oncogenic phenotypes and metabolism. We generated a panel of cell lines with increasing levels of fumarate accumulation by expressing FH variants with different catalytic efficiencies in HLRCC cells (Fig. 4). We observed a strong correlation between fumarate/malate ratios and proliferation rates (Supplementary Fig. S9D), which is consistent with a linear relationship between fumarate accumulation and tumorigenesis. For instance, while wildtype FH slowed proliferation of NCCFH1 cells, M151K had only a slight effect on the cell proliferation. As we better understand HLRCC cell transformation, it will be important to analyze how variants with partial activity might impact not just proliferation rates but also transformation, cell migration, and other aspects of tumorigenesis *in vivo*.

This panel of cell lines with different levels of fumarate accumulation also presented an opportunity to investigate how minor alterations to FH activity and fumarate levels affect HLRCC metabolism. TCA intermediates were among the most anticorrelated with fumarate levels; expression of wildtype FH increased total levels of citrate/isocitrate, cis-aconitate, and α-ketoglutarate without altering labeling patterns. Our findings demonstrate that glutamine is the main fuel source filling the TCA cycle and metabolized through reductive carboxylation, though there are differences between human and mouse FH/Fh1-deficient cells in metabolism of glutamine through reductive carboxylation^20,24,30,40^. Additionally, we observed a positive correlation between cystathionine and fumarate levels, which led us to investigate *de novo* cysteine and glutathione biosynthesis in HLRCC cells. We found that glutathione biosynthesis is increased in HLRCC cells, which may be a response to GSH-depletion occurring as a result of reactions with fumarate to form succinicGSH^27,28^. Thus, increased glutathione biosynthesis may help maintain redox balance in HLRCC cells. While our data provides a snapshot of fumarate-driven cellular metabolic rewiring *in vitro*, it will be important to determine whether similar metabolic changes occur as a result of fumarate accumulation *in vivo*.

In cells lacking FH there is an increase in adenylosuccinate, which has led others to predict that fumarate disrupts the synthesis of AMP^8,12,36^. Here we provide evidence that fumarate drives the reversal of ADSL in HLRCC cells, resulting in adenylosuccinate and SAICAR accumulation and broadly disrupting AMP and IMP biosynthesis (Fig. 5). We leveraged the different labeling patterns of aspartate (m+3) and fumarate (m+4) from U-^13^C-glutamine to analyze the incorporation of either substrate into adenylosuccinate and SAICAR. The observation of m+4 isotopomers in adenylosuccinate and SAICAR was consistent with reversal of ADSL in both reactions. Consistently, the increase in the AICAR/SAICAR ratio in cells expressing FH suggests that lowering fumarate restores the typical enzymatic activity of ADSL. Because *de novo* purine synthesis is diminished in cells lacking FH, we predicted that these cells rely on salvage from extracellular adenine and guanine to produce purine nucleotides. Consistent with this notion, we found that FH-deficient cells metabolize adenine and guanine to produce purine nucleotides. CRISPR-mediated genetic disruption of purine salvage pathway enzymes APRT and HGPRT reduced HLRCC cell proliferation and growth of xenograft tumors *in vivo*.

Targeting purine salvage using 6-MP, which competitively inhibits purine salvage pathway enzymes HGPRT and APRT, restricted HLRCC cell proliferation and the growth of xenograft tumors *in vivo*. Importantly, 6-MP is also metabolized to thioinosine monosphosphate (TIMP) and 6-methyl-thioinosine monophosphate (MeTIMP), which inhibit other aspects of purine biosynthesis including glutamine-5-phosphoribosylpyrophosphate amidotransferase, adenylosuccinate synthase, and IMP dehydrogenase (Supplementary Fig. S18D)^39^. Though our LC-MS methods were not able to detect 6-MP, TIMP, or MeTIMP in xenograft tumors, we were able to detect thiouric acid, which is an inactive metabolite of 6-MP, suggesting that 6-MP was taken up by the HLRCC xenograft tumors (Supplementary Fig. S18E). Further, tumors treated with 6-MP had decreased levels of purine nucleotides and intermediates.

It is likely that fumarate accumulation renders cells more sensitive to 6-MP due to: 1) a necessary reliance on the purine salvage pathway resulting from the fumarate-driven reversal of ADSL enzymatic activity, and/or 2) increased nucleoside uptake and metabolism, including 6-MP. We expect that inhibitors that specifically target the purine salvage pathway without affecting other aspects of purine biosynthesis would be effective at restricting the growth of FH-deficient cells with minimal effect on non-transformed, FH-expressing cells. Re-purposing fairly well-tolerated agents like 6-MP or azathioprine could be an exciting pathway to rapidly translate these discoveries to the clinic.

Together our findings highlight the need to establish cancer-associated risks for germline *FH* alterations, suggest the pathogenicity of many patient-associated *FH* variants, and demonstrate that fumarate accumulation disrupts purine biosynthesis in HLRCC cells which may be targetable using existing clinical compounds.

## Methods

### Ethical regulations

The use of tissue and clinical information was permitted through our institutional review board (IRB) approved protocol 19-001174. Interrogation of patient-derived *FH* variants found after genetic risk assessment was permitted through our IRB approved protocol #21-000887.

All animal experiments were approved by the UCLA Animal Research Committee (ARC), and we complied with all relevant ethical regulations. Mice were housed in a pathogen-free facility at University of California, Los Angeles (UCLA). We used the minimum number of mice necessary to obtain significant and reliable data.

### Bacteria-expression plasmid cloning

Gibson cloning (NEB) was used to generate a bacteria expression plasmid containing 6His-tagged FH, lacking the first 44 amino acids, corresponding to the mitochondrial targeting sequence. This resulting plasmid, pet28a-FH(45-510), was used to generate expression plasmids with FH variants using a QuikChange II kit (Agilent). Sanger sequencing was used to confirm sequence identity.

### Recombinant FH and cell-free assays

BL21-STAR *E. coli* cells carrying pet28a-FH(45-510) or mutant constructs were grown to density in Luria Broth with kanamycin (50 µg/mL) by shaking overnight at 37°C with agitation. The culture was diluted 20-fold then allowed to grow for 2 hours at 37°C with agitation. IPTG was added to 0.5 mM then incubated at 37°C for 2 hours with agitation. *E. coli* was harvested by centrifugation, then resuspended in bacteria lysis buffer (50 mM sodium phosphate pH 7.4, 300 mM NaCl, 1% triton X-100, 10 mM imidazole, 20% glycerol, 1 mM TCEP, protease inhibitors, and 5 mg/mL lysozyme). Bacteria were lysed by 6 rounds of freeze-thaw, then the cell extract was cleared by centrifugation. His-tagged FH(44-511) was purified from the lysate using HisPur^TM^ Cobalt Spin Columns (Thermo Fisher). Columns were washed using with buffer A (50 mM sodium phosphate pH 7.4, 300 mM NaCl, 0.01% Tween-20, and 10 mM imidazole) and recombinant protein was eluted using buffer B (50 mM sodium phosphate pH 7.4, 300 mM NaCl, 0.01% Tween-20, and 300 mM imidazole). The eluate was dialyzed against buffer C (50 mM sodium phosphate pH 7.4, 300 mM NaCl, and 1 mM TCEP) overnight at 4 °C. The protein concentration was quantified using Bradford reagent then diluted to 10 µM in 50% glycerol, 25 mM sodium phosphate pH 7.4, 150 mM NaCl, and 0.5 mM TCEP. Recombinant was stored at −20°C with no detectable decrease in activity over several months. To validate purity, recombinant proteins were resolved on SDS-Polyacrylamide gel electrophoresis followed by Coomassie staining.

### Cell-free assays

The activity of recombinant FH was determined by quantifying the change in absorbance at 250 nm a wavelength which is absorbed by fumarate but not by malate. Recombinant FH was diluted to 10 nM in reaction buffer (50 mM sodium phosphate pH 7.4, 150 mM NaCl, and 0.25% tween-20). Reactions were started by adding fumarate or malate and various concentrations, following which absorbance was measured every minute for 30 minutes on an Agilent BioTek Synergy H1. All reactions were carried out at 37°C. For all plates, a standard curve was measured in technical triplicates in order to quantify fumarate depletion (forward reaction) or fumarate generation (reverse reaction). A linear regression was used for to model reaction rates. Rates were fitted to a Michaelis-Menten model and associated V_max_, K_m_, and k_cat_ were calculated with standard errors using GraphPad Prism (RRID:SCR_002798). Catalytic efficiencies were calculated as k_cat_/K_m_.

### Native-PAGE

Multimerization status was analyzed by resolving proteins on precast Mini-PROTEAN® TGX^TM^ gels (BioRad) using 1X Tris/Glycine buffer (BioRad) with 0.02% Coomassie G in the cathode buffer. NativeMark^TM^ Unstained Protein Standard (Thermo Fisher) was used to validate determine the molecular weight associated with each band. Bands were quantified using ImageJ.

### Cell lines, cell culture and reagents

UOK-262 (RRID:CVCL_1D72) cells were provided from the laboratory of Brian Shuch (UCLA). The NCCFH1 cell line is owned by the National Cancer Centre of Singapore Pte Ltd. The patient-derived xenograft line xp-152 was provided from Xiankai Sun (University of Texas, Southwestern). The cell line xp-152-cl was generated by dissociating a 3 mm^3^ portion of xp-152, then passaging adherent cells for > 40 passages such that 1) surviving cells appeared to be immortal, 2) we observed no passage-to-passage variation in proliferation rates, and 3) we detected no fumarate oxidation. All HLRCC cell lines were grown in RPMI supplemented with 10% FBS, 1% penicillin/streptomycin, and 2mM pyruvate. HEK293T cells were obtained from American Type Culture Collection (ATCC, RRID:CVCL_0063) and cultured in DMEM supplemented with 10% FBS and 1% penicillin/streptomycin. All cells were maintained at 37°C with 5% CO_2_, and monitored regularly for mycoplasma using either a MycoAlert detection kit (Lonza, LT07-318) or by PCR analysis using an established protocol^41,42^.

### Lentiviral plasmid cloning

Standard PCR/restriction enzyme cloning was used to produce a mammalian lentiviral expression plasmid containing FH with a c-terminal V5-tag. The resulting plasmid, pLenti6-FH-V5, was used as a template for generating expression plasmids with *FH* variants using the QuikChange II kit (Agilent). Sanger sequencing was used to confirm sequence identity for all plasmids. The control plasmid, pLenti6-GFP, was a gift from Daniel Haber (Addgene plasmid # 35637; http://n2t.net/addgene:35637; RRID:Addgene_35637)^43^.

### Lentivirus production and transduction

HEK293T cells were transfected with a pLenti6 transfer plasmid, pRSV-Rev, pMDLg/pRRE, and pCMV-VSV-G. pRSV-Rev was a gift from Didier Trono (Addgene plasmid # 12253; http://n2t.net/addgene:12253; RRID:Addgene_12253). pMDLg/pRRE was a gift from Didier Trono (Addgene plasmid # 12251; http://n2t.net/addgene:12251; RRID:Addgene_12251). pCMV-VSV-G was a gift from Bob Weinberg (Addgene plasmid # 8454; http://n2t.net/addgene:8454; RRID:Addgene_8454). 48 hours post-transfection media was collected and filtered at 0.2 µm, then immediately used to transduce HLRCC cells.

HLRCC cells were transduced with lentivirus containing 4 µg/ml polybrene. 16 hours after transduction, media containing lentivirus was replaced with fresh media. 24 hours later, media was replaced with fresh media containing 2 µg/ml blasticidin.

### Cell proliferation assays

Cells were seeded at 50,000 cells per well of a 6-well dish in triplicate. 24 hours later, three wells were counted using a particle counter. For remaining wells, cells were washed twice with PBS and media was replace, then 48 hours later all remaining wells were counted. Cell counts were used to model the growth grates (k) using the exponential growth equation.

### LC-MS and analysis

Cells were seeded in triplicate in 6-well plates such that at 16-24 hours, cells were at 70-90% confluency. For ^13^C-, ^2^H, or ^15^N-labeling experiments, cells were washed twice with PBS then treated for 6 hours with DMEM containing 10% dialyzed-FBS, 1% penicillin/streptomycin, and tracers. The following tracers were used in place of unlabeled metabolites in the media: 25 mM U-^13^C-glucose, 4 mM U-^13^C-glutamine, 0.2 mM 3,3,3’,3’-^2^H-cystine, 4 mM amide-^15^N-glutamine, 50 µM 8-13C-adenine, and 50 µM 8-13C-guanine (all from Cambridge Isotope Laboratories).

Cells were washed twice with cold ammonium-acetate, pH 7.4, then 80% methanol containing 10 nM trifluoromethanesulfonate (internal standard) was added and cells were incubated at −80°C for 15 minutes. Cells were scraped, transferred to microcentrifuge tubes, vortexed well, then cleared by centrifugation. Supernatant was transferred to a new tube and dried over a continuous flow of N_2_ for 1 hour. Metabolites were maintained at −80°C until analyzed by LC-MS.

Dried metabolites were reconstituted in 100 µl of 50% acetonitrile. Samples were vortexed and cleared by centrifugation. 10 µl of of the metabolite mixture was injected per analysis. Samples were run on a Vanquish (Thermo Scientific) UHPLC system with mobile phase A (20 mM ammonium carbonate, pH 9.7) and mobile phase B (100% acetonitrile) at a flow rate of 150 µL/min on a SeQuant ZIC-pHILIC Polymeric column (2.1 × 150 mm 5 μm, EMD Millipore) at 35°C. Separation was achieved in three steps: 1) a linear gradient from 80% B to 20% B for 20 minutes, 2) a linear gradient from 20% B to 80% B for 0.5 minutes, and 3) 80% B for 7.5 minutes. The UHPLC was coupled to a Q-Exactive (Thermo Scientific) mass analyzer running in polarity switching mode.

Each file was centroided and converted into positive and negative mzXML files using msconvert from ProteoWizard. MZmine 2 was used to generate ion chromatograms from MS1 spectra via a built-in Automated Data Analysis Pipeline (ADAP) chromatogram module^44^. Peaks were detected via the ADAP wavelets algorithm. Peaks were aligned across all samples via the Random sample consensus aligner module, gap-filled, and assigned identities using an exact mass MS1(+/-15ppm) and retention time RT (+/-0.5min) search of our in-house MS1-RT database. Our MS1-RT database consists of 164 metabolites for which we have analyzed standards. Peak boundaries and identifications were then further refined by manual curation.

Peaks were quantified by area under the curve integration. Fumarate/malate and other ratios were calculated by dividing peak areas. If stable isotope tracing was used in the experiment, the peak areas were additionally processed via the R package AccuCor 2 to correct for natural isotope abundance^45^. Peak areas for each sample were normalized by the measured area of the internal standard trifluoromethanesulfonate (present in the extraction buffer) and by the number of cells present in the extracted well. MetaboAnalyst 5.0 (RRID:SCR_015539) was used to conduct over-representation metabolite set enrichment analysis using metabolic pathways annotated by KEGG^46,47^.

### Immunohistochemistry

Immunostaining for 2SC was performed on formalin-fixed, paraffin-embedded tissue (FFPE) sections using rabbit anti-human 2SC polyclonal antibody (Cambridge Research Biochemicals Cat# crb2005017, RRID:AB_2892588) at a dilution of 1:1000 with a modified Agilent FLEX Envision detection system with an Automated Autostainer (AS48Link). All FFPE slides were pretreated with Heat Induced Epitope Retrieval (HIER) in Dako PTLink using Envision FLEX Target Retrieval solution and incubated 97℃ for 20 minutes.

### Immunoblotting

Cells were lysed in a buffer containing 40 mM HEPES (pH 7.4), 120 mM NaCl, 1 mM EDTA, 1% NP40, 1 mM DTT, 1 mM sodium orthovanadate, 20 mM sodium flouride, 10 mM β-glycerophosphate, 10 mM sodium pyrophosphate, 2 µg/ml aprotinin, 2 µg/ml leupeptin, 2 µg/ml pepstatin, and 1 mM PMSF. Immunoblotting was performed using standard procedures. The following commercially available antibodies were used: GFP (Cell Signaling Technology Cat# 2956, RRID:AB_1196615; dilution 1:1000), FH (Cell Signaling Technology Cat# 4567, RRID:AB_11178522; dilution 1:1000), v5-HRP (Thermo Fisher Scientific Cat# R961-25, RRID:AB_2556565; dilution 1:50,000), and α-tubulin (Sigma-Aldrich Cat# T6074, RRID:AB_477582; dilution 1:10,000).

### Gene expression

RNA-sequencing data from HLRCC-associated primary kidney tumors, metastases, and adjacent normal kidney tissue was obtained from NCBI Gene Expression Omnibus – GSE-157256^48^. GEO RNA-seq Experiments Interactive Navigator (GREIN) was used to obtain counts per million for all samples^49^.

### Mouse cell line xenografts

5E6 NCCFH1 or xp-152-cl cells in a 1:1 mixture of Matrigel and PBS were injected subcutaneously into the flanks of 7 week-old NSG mice. Tumors were monitored regularly for growth by calipering, and tumor volume was calculated as (*L*x*W*^2^)/2 where *L* is length and *W* is width. Every experimental group contained an equal number of male and female mice. For 6-MP experiments, treatment began when the tumors reach 150 mm^3^. 6-MP was resuspended in DMSO at a high concentration, which was then diluted into injection buffer (0.9% NaCl, 50 mM glycine, pH 10), which was then delivered to each mouse via intraperitoneal injection every other day. Each mouse received no more than 0.5 µL/g DMSO per dosing. Tumors were harvested when the first tumor reached our designated endpoint (1000 mm^3^) or mice started to show signs of distress. Tumors were snap frozen in liquid nitrogen, and serum was collected and stored at −80 °C.

### Cell cycle and apoptosis analysis

Cells were seeded at a low density such that following treatment, cells would be sub-confluent for the cell cycle and apoptosis analysis. Cells were treated for 48 hours with media containing 6-MP or DMSO. A control apoptosis group recieved 0.1 µM staurosporine for 24 hours. Cell suspensions were co-stained with Vybrant^TM^ DyeCycle^TM^ Green (Thermo Fisher) and Annexin V, Alexa Fluor 647 conjugate (Thermo Fisher) for 30 minutes at room temperature in conjugation buffer (10 mM HEPES, 140 mM NaCl, and 2.5 mM CaCl_2_, pH 7.4). Cells were analyzed on a BD LSRFortessa^TM^ cell analyzer.

### Replication

All experiments were performed with a minimum of three biological replicates for each experiment. All mouse experimental groups included ≥8 mice, with equal numbers of male and female mice.

### Quantification and statistical analyses

Graphs represent the mean ± standard error of the mean (SEM). For analyses with two or more groups, a two-way ANOVA was used to generate p-values and percent variation. For analyses with only one group, a two-tailed Students t-test was used to generate p-values. p-values are represented as *p ≤ 0.05; **p ≤ 0.01; ***p ≤ 0.001; ****p ≤ 0.0001. GraphPad Prism was used to model Michaelis-Menten kinetics and determine the fitted variables with standard errors. For Native-PAGE and densitometry experiments, ImageJ (RRID:SCR_003070) was used to quantify band density. The R package stats (version 3.6.2) was used to perform principal component analysis. BioRender.com was used to create visualizations, UCSF Chimera was used to visualize crystal structures, and GraphPad Prism was used to create graphs.

---

**Supplementary Fig. S1:**
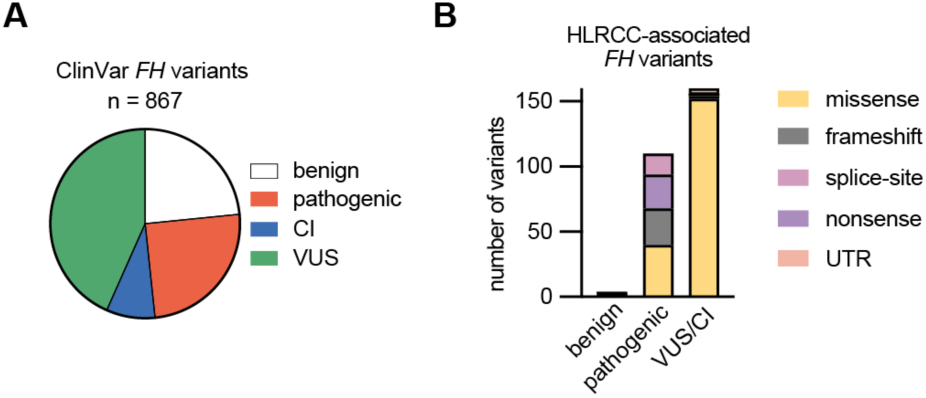
Analysis of *FH* variants and their associated interpretations of pathogenicity. (**A**) Pie chart showing proportion of all *FH* variant annotated in ClinVar and their classification as either benign (or likely-benign), pathogenic (or likely-pathogenic), CI, or VUS. (**B**) HLRCC-associated, protein-altering *FH* variants split into benign (or likely-benign), pathogenic (or likely-pathogenic), and VUS/CI. Colors represent the type of mutation, showing that nearly all of the VUS/CI are missense mutations.

**Supplementary Fig. S2:**
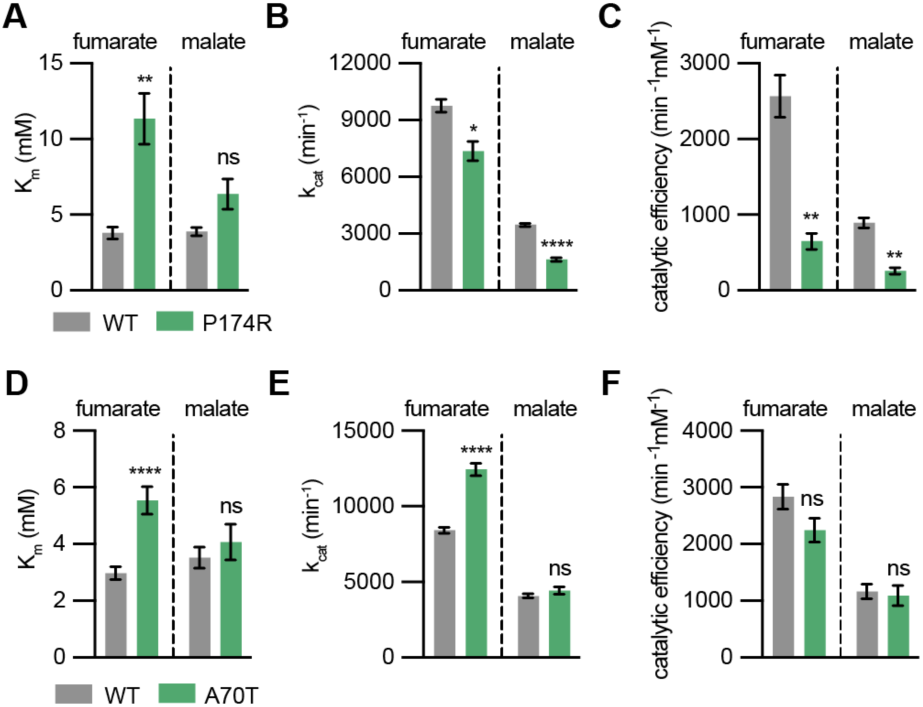
Kinetic parameters for active/partially-active VUS/CI. (**A**) K_m_, (**B**) k_cat_, and (**C**) catalytic efficiency for wildtype (WT) and P174R. Data is reported for fumarate (left) and malate (right) indicating compromised enzymatic activity in both directions. (**D**) K_m_, (**E**) k_cat_, and (**F**) catalytic efficiency for WT and A70T. Data is reported for fumarate (left) and malate (right). While K_m_ and k_cat_ for fumarate are higher for A70T, catalytic efficiencies are nearly identical to WT, suggesting that this variant is comparable in activity to WT enzyme.

**Supplementary Fig. S3:**
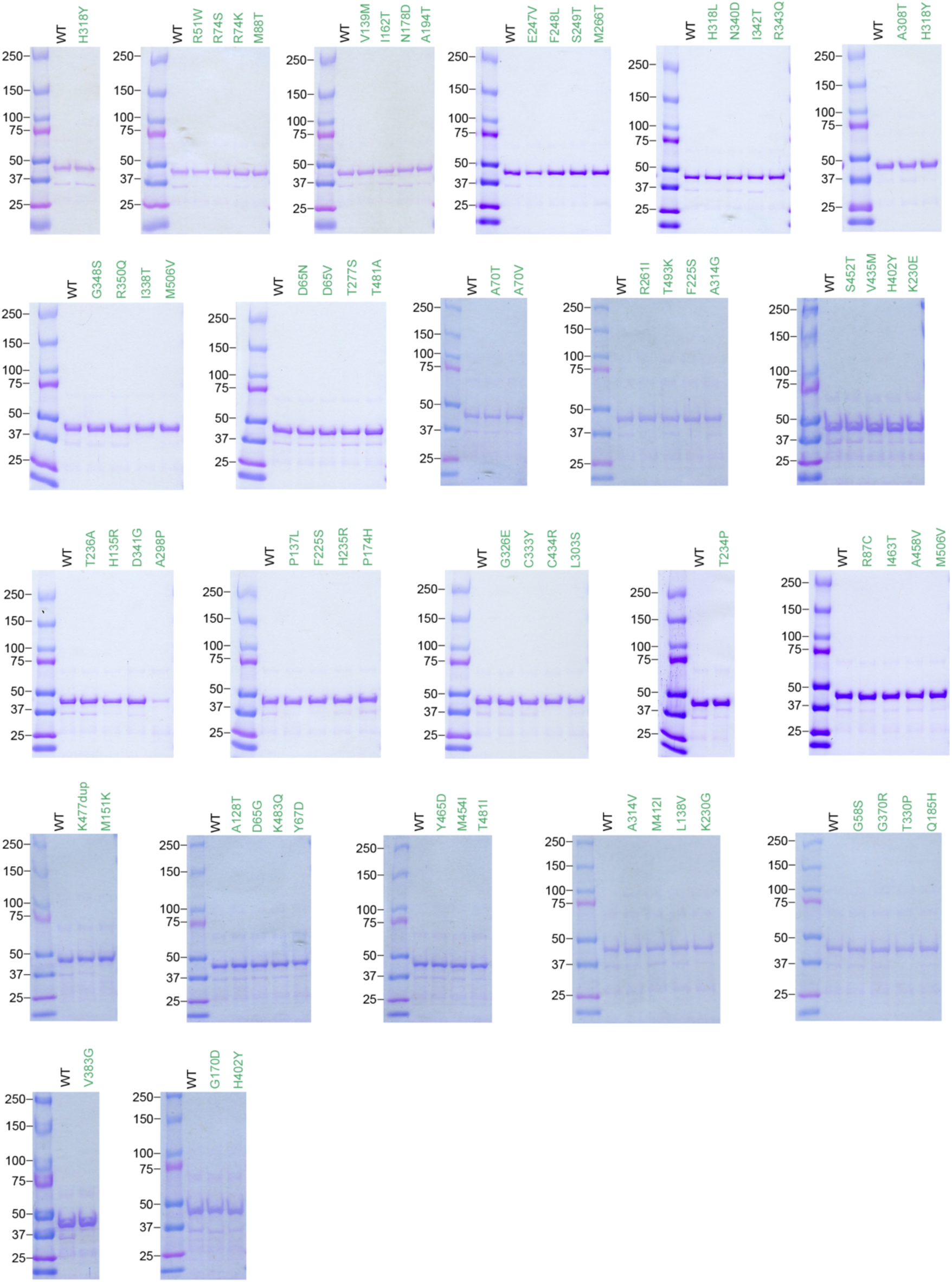
Recombinant FH produced for 74 VUS/CI. Recombinant FH was produced and purified from *E. coli* as shown by SDS-PAGE gels stained by Coomassie.

**Supplementary Fig. S4:**
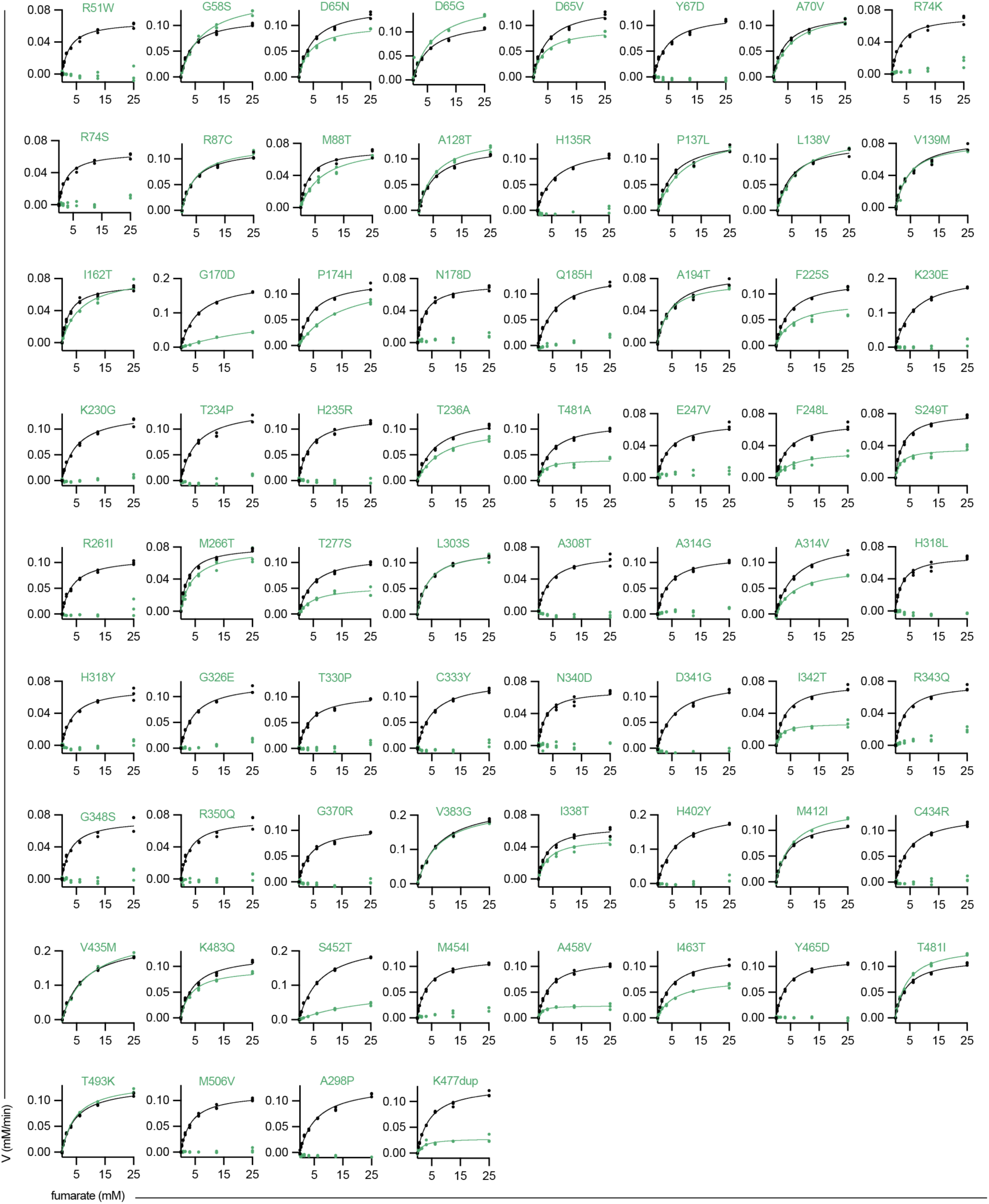
Michaelis-Menten graphs for VUS/CI incubated with fumarate. Recombinant FH was incubated with increasing concentrations of substrate fumarate and conversion to product was measured. Wildtype (WT) FH was run simultaneously for each variant, as indicated by the black symbols/lines. VUS/CI have a range of enzymatic activities.

**Supplementary Fig. S5:**
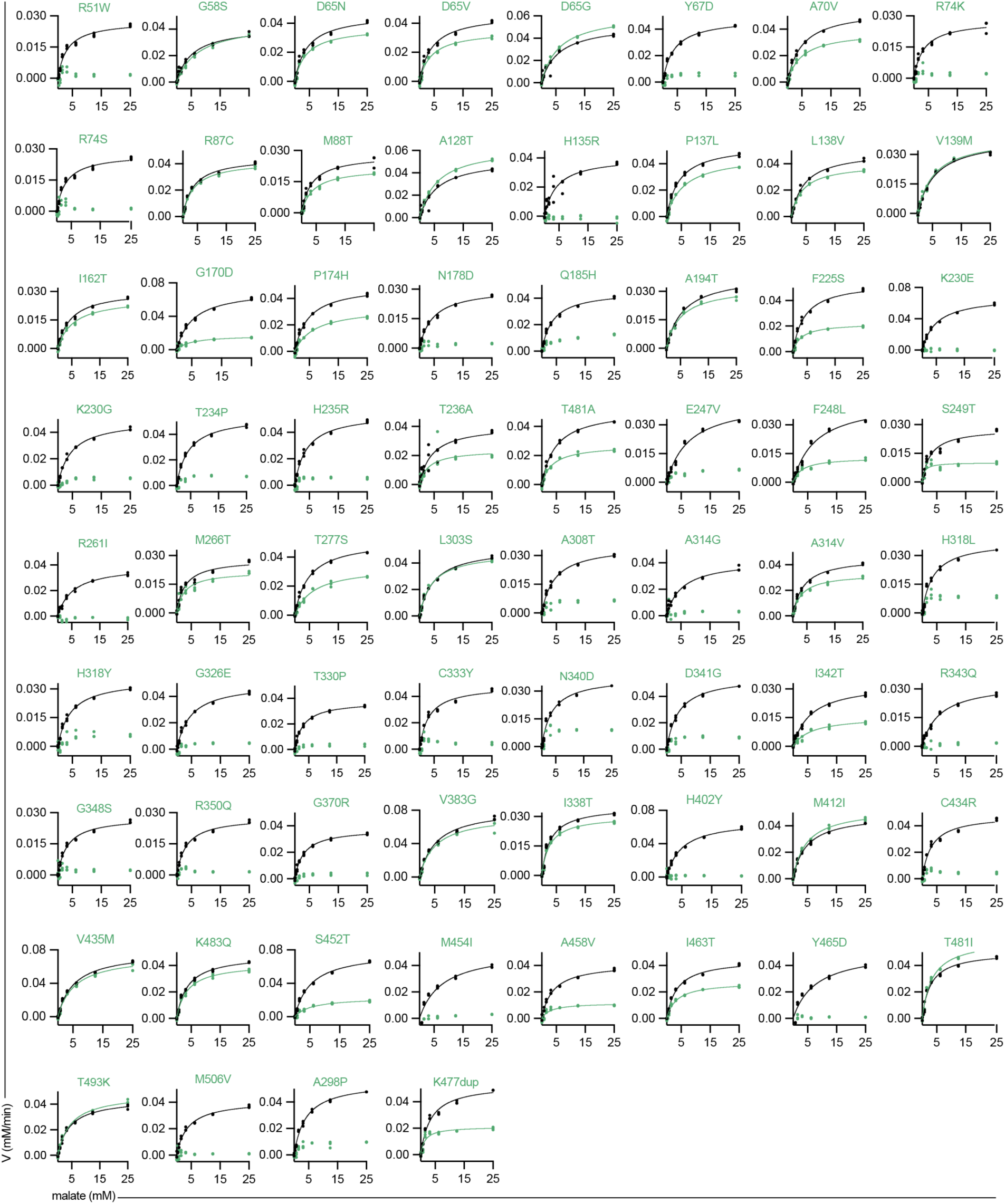
Michaelis-Menten graphs for VUS/CI incubated with malate. Recombinant FH was incubated with increasing concentrations of substrate malate and conversion to product was measured. Wildtype (WT) FH was run simultaneously for each variant, as indicated by the black symbols/lines. VUS/CI have a range of enzymatic activities.

**Supplementary Fig. S6:**
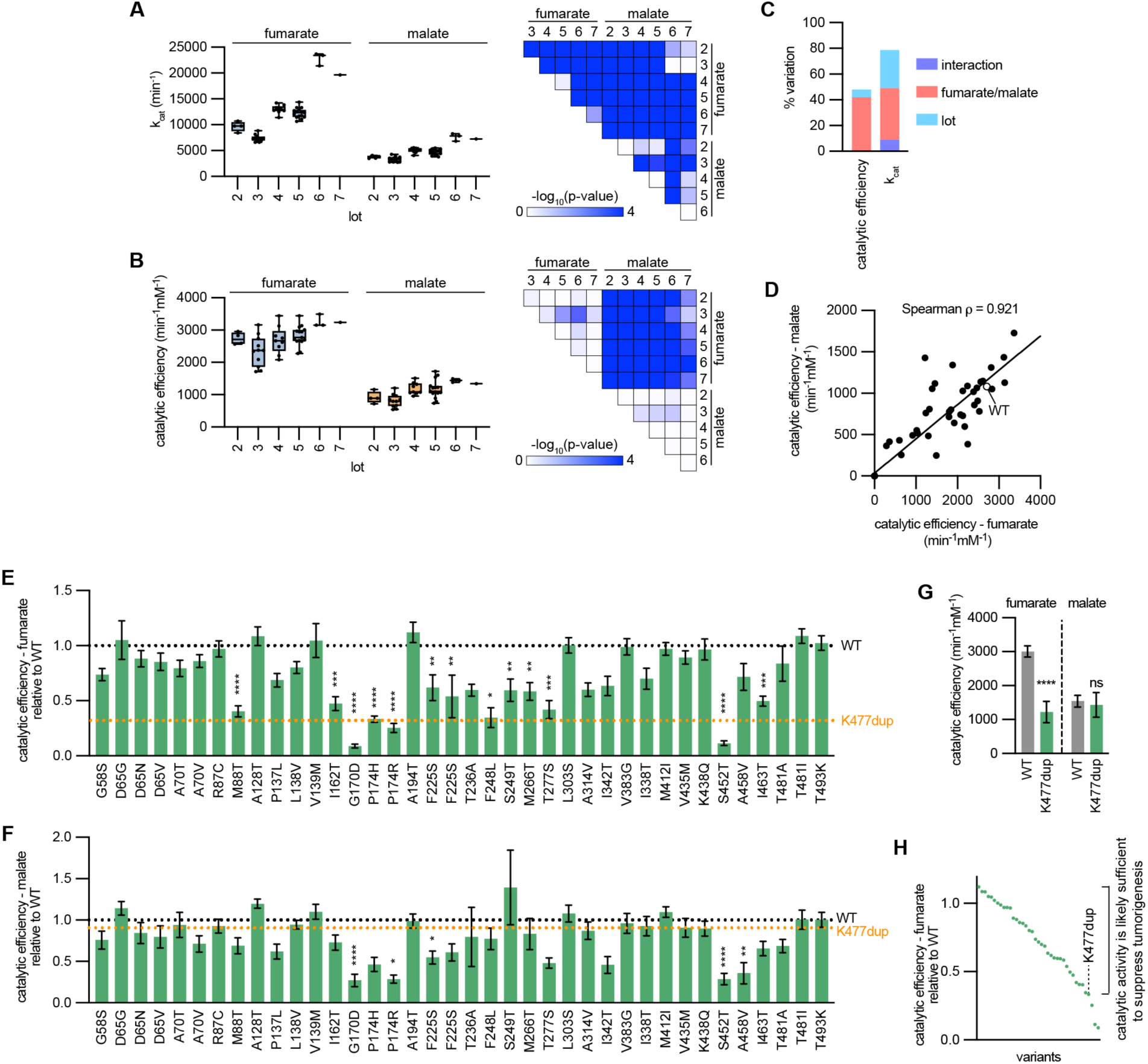
Broad analysis of recombinant FH. (**A**) k_cat_ for wildtype (WT) FH. All analyses (3 reaction series per plate) are reported as symbols, broken down into lots and substrates. For all possible comparisons, −log_10_(p-value) is reported as a heatmap, showing widespread change across lots and substrates. (**B**) Catalytic efficiencies for WT FH, broken down into lots and substrates. −log_10_(p-value) is reported as a heatmap, demonstrating reduced variability across lots, but maintaining variability between substrates. (**C**) A two-way ANOVA was performed on all WT data reported in panels a and b. The protein lot had a greater contribution to the total dataset variation for k_cat_ than for catalytic efficiency. For catalytic efficiency, substrate had >4-times the contribution to the total variation in the dataset than the lot. (**D**) Linear regression of catalytic efficiencies for fumarate and malate. The data correlated well (Spearman ρ = 0.921), suggesting that mutations typically alter catalytic activity for both directions. Relative catalytic efficiency for (**E**) fumarate and (**F**) malate for all variants with measurable catalytic activity. (**G**) Catalytic efficiencies for K477dup. (**H**) Relative catalytic efficiencies for all variants with measurable catalytic activity. K477dup is indicated by a dashed line, which might serve as a benchmark, where variants with higher catalytic efficiencies could be sufficient to suppress tumorigenesis.

**Supplementary Fig. S7:**
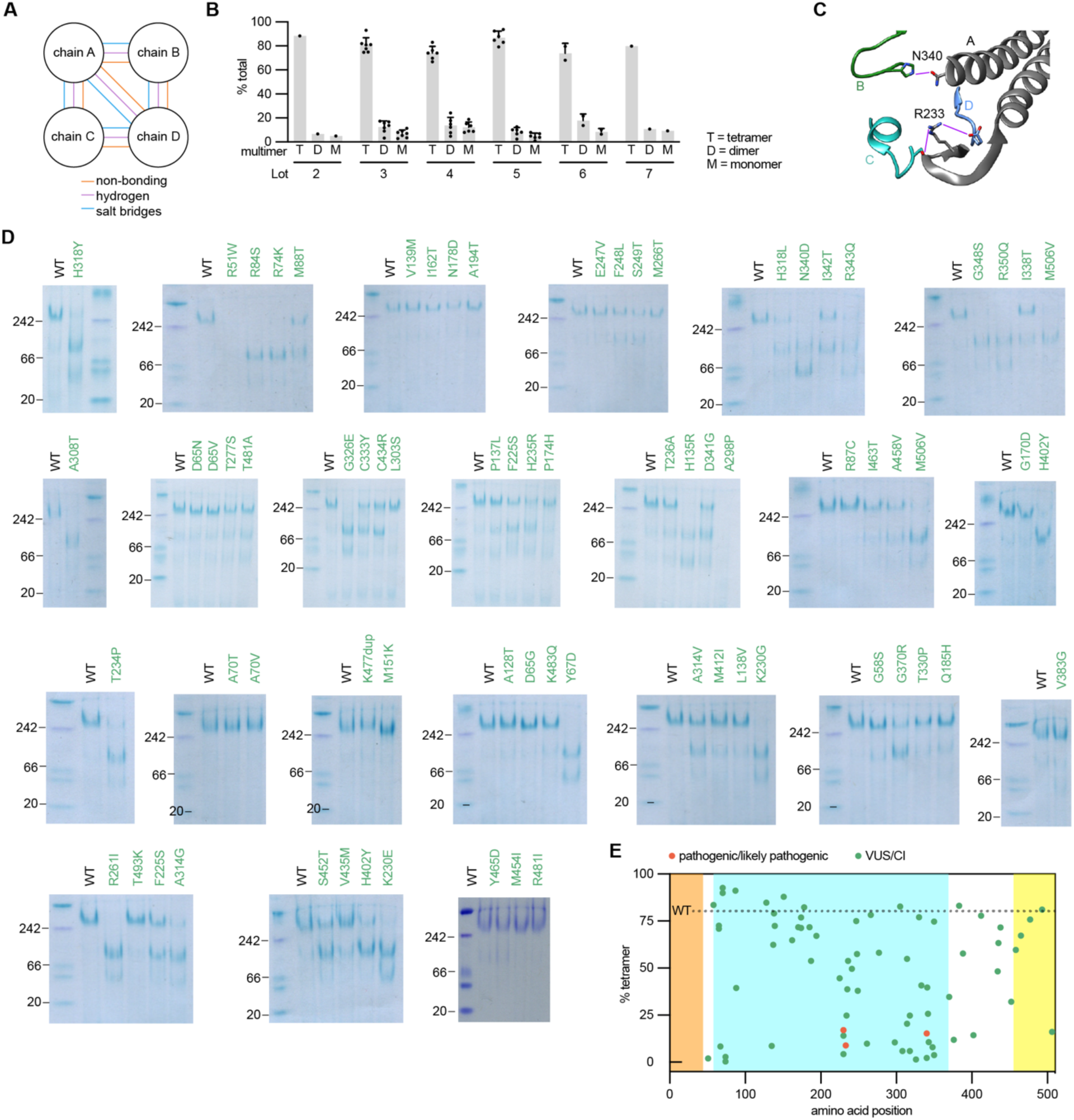
Multimerization status of 74 VUS/CI. (**A**) Schematic demonstrating the binding pattern of multimers in tetrameric FH. Residues interact with other chains as non-bonding, hydrogen bonding, and salt bridge-forming. (**B**) Analysis of wildtype (WT) FH for each gel shows little variation in multimerization patterns across lots. (**C**) Crystal structure of FH (3E04) with R233 and N340 and their hydrogen bonds with residues in other chains. (**D**) Native-PAGE gels stained with Coomassie showing variation in multimerization status. (**E**) Graph indicating the percent tetramerization for 74 FH variants. WT percent tetramerization is indicated as a dotted line.

**Supplementary Fig. S8:**
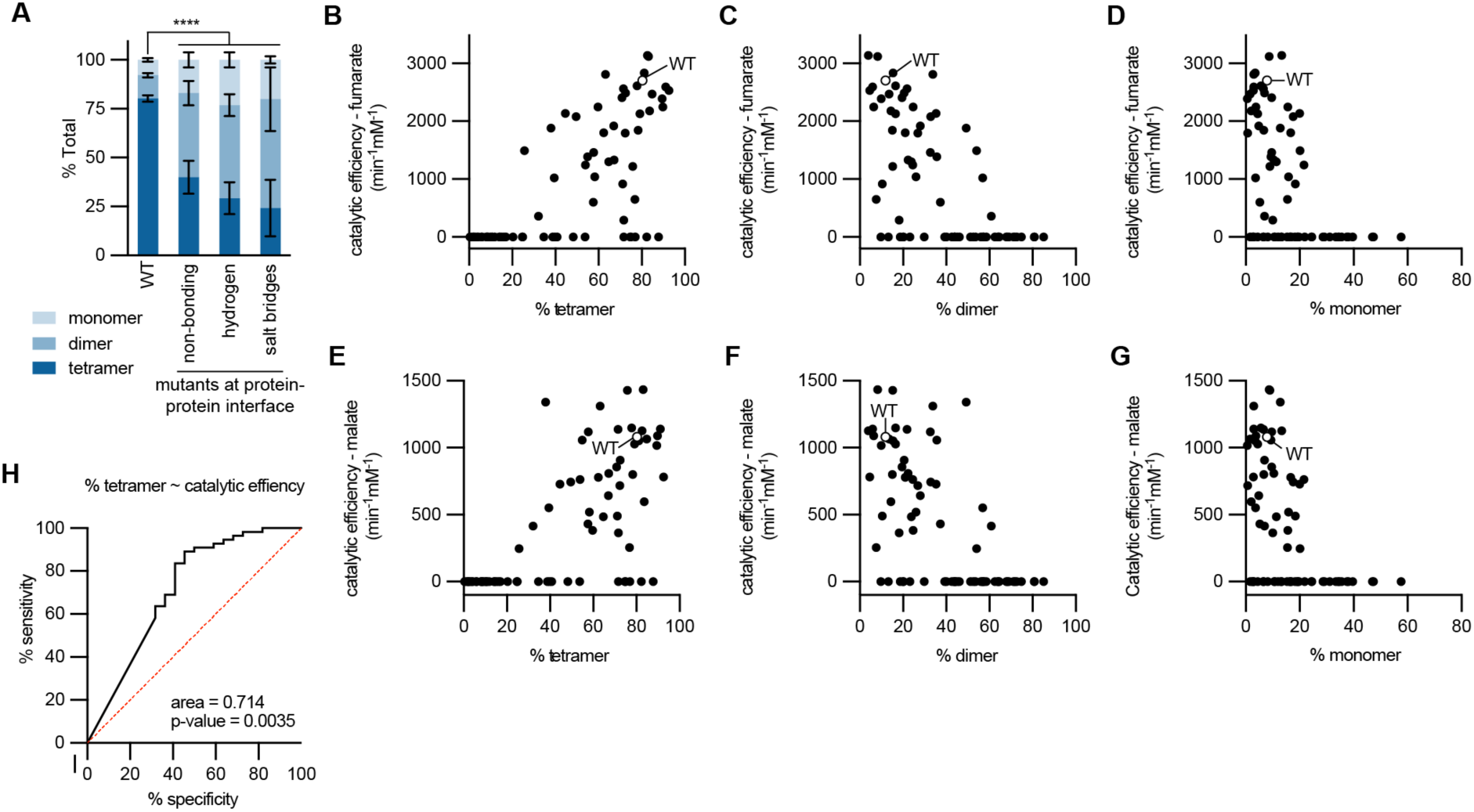
Tetramerization correlates with catalytic activity. (**A**) Percent of multimerization species for wildtype (WT) and variants that occur at multimerization interfaces, including those that occur at residues participating in non-bonding, hydrogen bonding, and salt bridge-forming interactions. Catalytic efficiencies for fumarate compared to the percent (**B**) tetramer, (**C**) dimer, and (**D**) monomer. Catalytic efficiencies for malate compared to the percent (**E**) tetramer, (**F**) dimer, and (**G**) monomer. The positive correlation between catalytic efficiency and percent tetramerization suggests that tetramerization is necessary for catalytic activity. (**H**) Receiver operating characteristic (ROC) curve investigating the diagnostic ability of % tetramerization for predicting catalytic activity, suggests that multimerization status is weakly predictive of enzyme activity.

**Supplementary Fig. S9:**
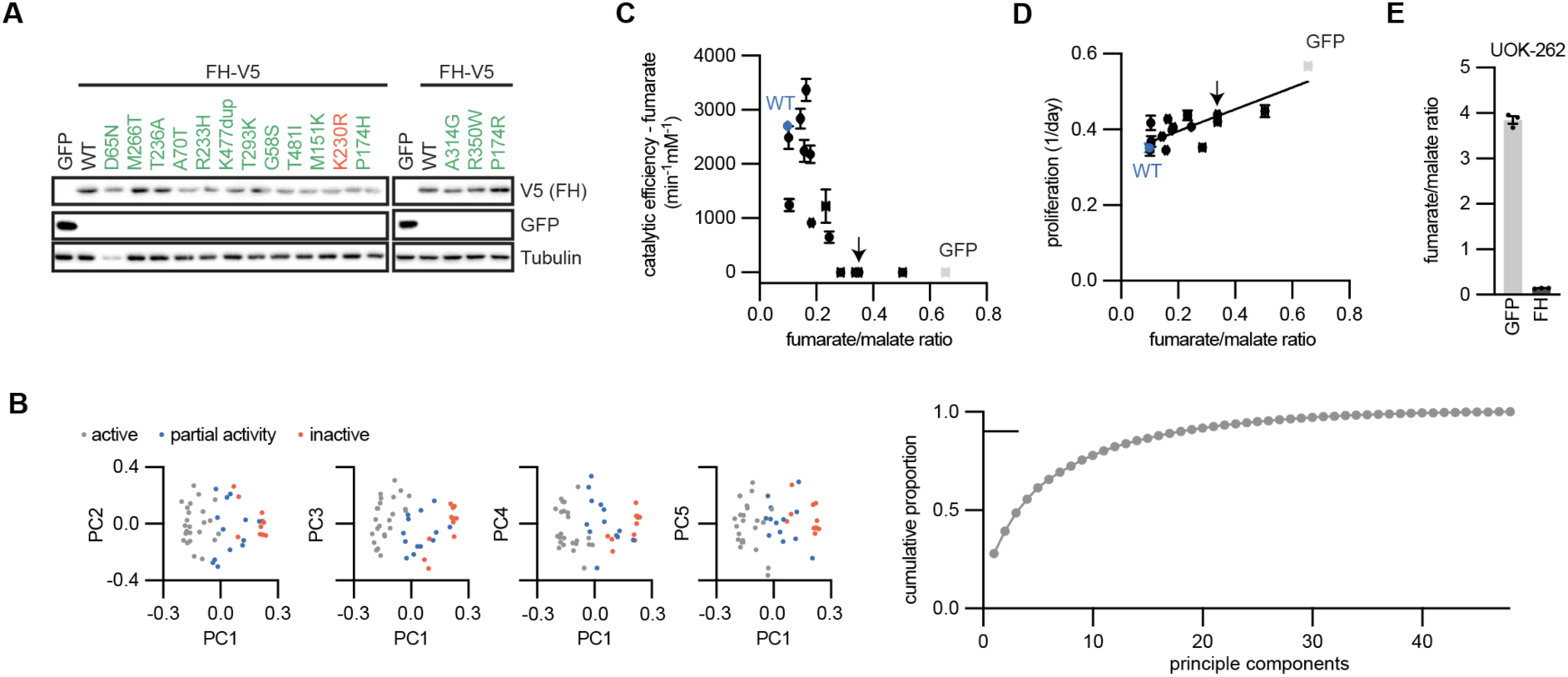
Expression of FH variants in HLRCC cells. Exogenous expression of GFP, V5-tagged FH, and V5-tagged FH variants in NCCFH1 cells. (**A**) Immunoblot showing expression of transgenes. (**B**) Principal component analysis using targeted metabolomics data. Data for the first 5 principal components are colored by expression of active, partially active, or inactive variants. Samples cluster together according to variant activity status. Also shown is the proportion of explained variance for principal components reaches 0.9 at PC18. (**C**) Catalytic efficiency as calculated in the cell-free assays correlates with fumarate/malate ratios in HLRCC cells expressing the corresponding variant. (**D**) Proliferation rates correlated with fumarate/malate ratios in HLRCC cells expressing the corresponding variants. (**E**) Fumarate/malate ratios UOK-262 cells expressing FH compared to control (GFP) cells.

**Supplementary Fig. S10:**
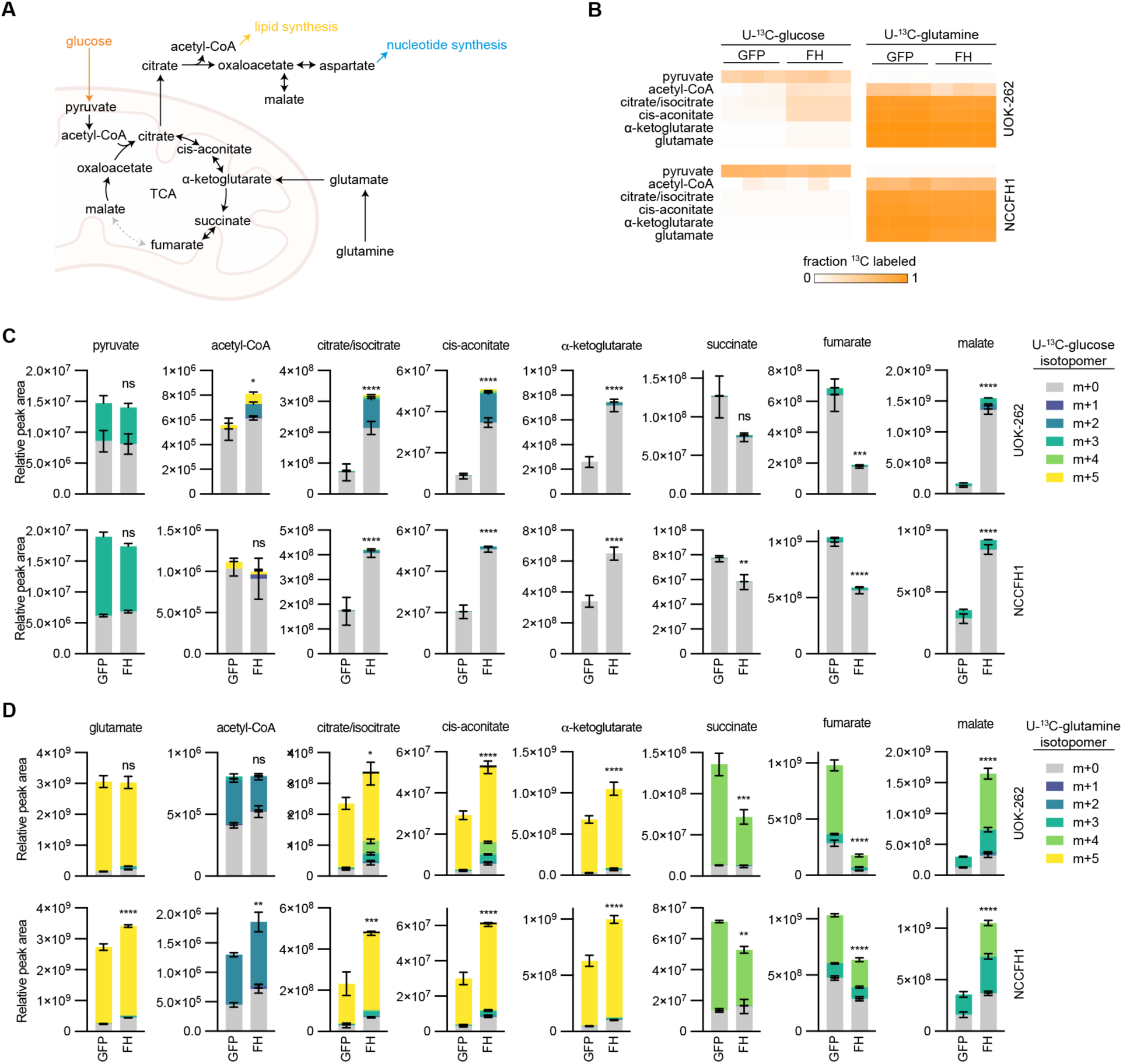
Fumarate accumulation decreases TCA intermediates. (**A**) Schematic showing the oxidative decarboxylation and reductive carboxylation as possible routes for glutamine and glucose metabolism in the TCA cycle. (**B**) Heatmap showing the fraction of ^13^C-labling for metabolites. The labeled atoms originated from U-^13^C-glucose (left) or U-^13^C-glutamine (right). (**C**) Relative metabolite levels for pyruvate and TCA intermediates in HLRCC cells expressing GFP or FH and treated with U-^13^C-glucose. Bars are colored by the relative abundance of isotopomers. (**D**) Relative metabolite levels for pyruvate and TCA intermediates in HLRCC cells expressing GFP or FH and treated with U-^13^C-glutamine. Bars are colored by the relative abundance of isotopomers.

**Supplementary Fig. S11:**
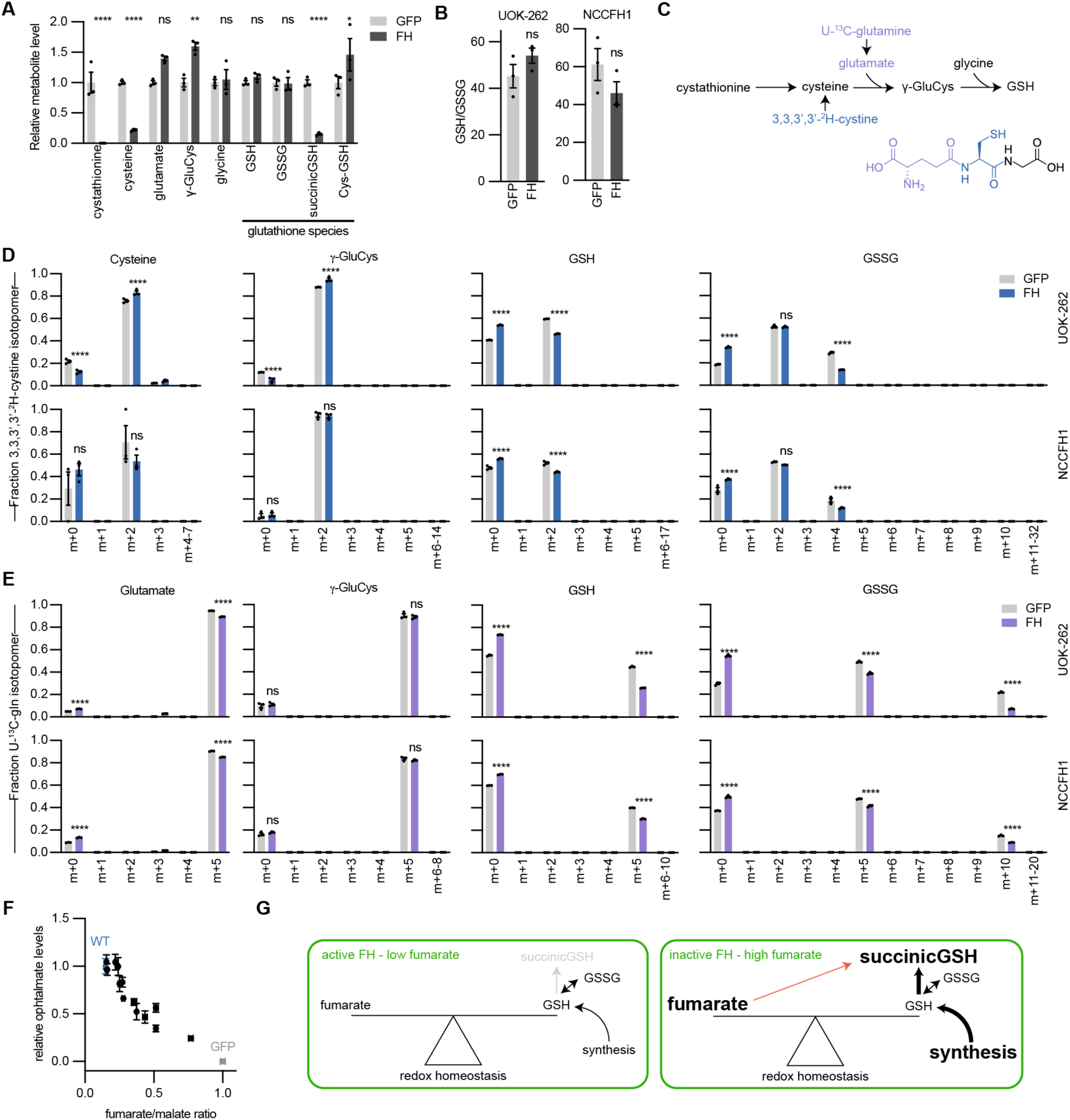
FH loss enhances de novo glutathione biosynthesis. (**A**) Relative levels of intermediates of de novo glutathione biosynthesis. (**B**) Ratio of reduced glutathione to oxidize glutathione for HLRCC cell lines expressing GFP or FH. (**C**) Schematic illustrating *de novo* glutathione biosynthesis and colored by contribution glutamine and cystine. HLRCC cells expressing GFP or FH were treated with U-^13^C-glutamine and 3,3,3,3-^2^H-cystine and isotopomers of glutathione biosynthesis intermediates were quantified. (**D**) Cysteine and γ-glutamylcysteine (γ-GluCys) were almost completely labeled from 3,3,3,3-^2^H-cystine. GSH and GSSG had a higher degree of labeling from 3,3,3,3-^2^H-cystine. (**E**) Glutamate and γ-glutamylcysteine (γ-GluCys) were almost completely labeled from U-^13^C-glutamine. GSH and GSSG had a higher degree of labeling from 3,3,3,3-^2^H-cystine. (**F**) fumarate/malate ratio most negatively correlated with ophthalmate, a glutathione analogue and biomarker of reduced oxidative stress. (**G**) Schematic illustrating our proposed model for how fumarate and glutathione biosynthesis are coordinated to maintain oxidative homeostasis. When FH activity is lost, fumarate accumulates, which reacts with GSH to produce succinicGSH. As a result, glutathione biosynthesis is increased.

**Supplementary Fig. S12:**
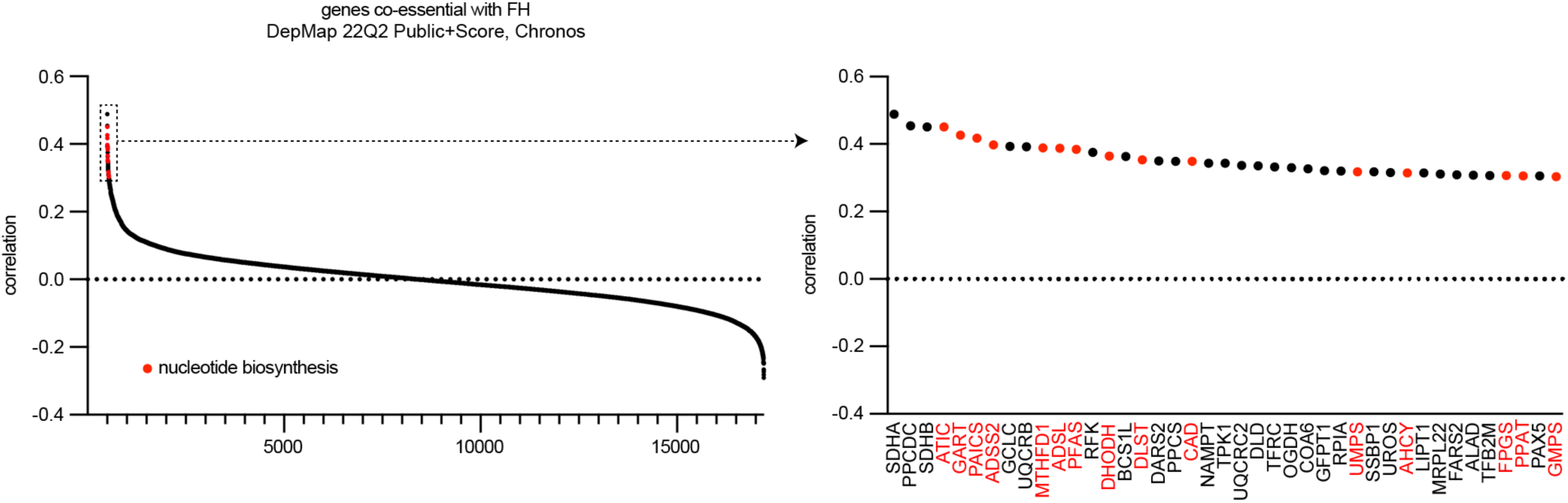
Coessentiality of FH and nucleotide biosynthesis genes. Correlation of DepMap 22Q2 essentiality scores for FH compared to all other genes. Nucleotide biosynthesis genes are indicated as red dots. Top 41 genes from the co-essentiality analysis are shown on the right.

**Supplementary Fig. S13:**
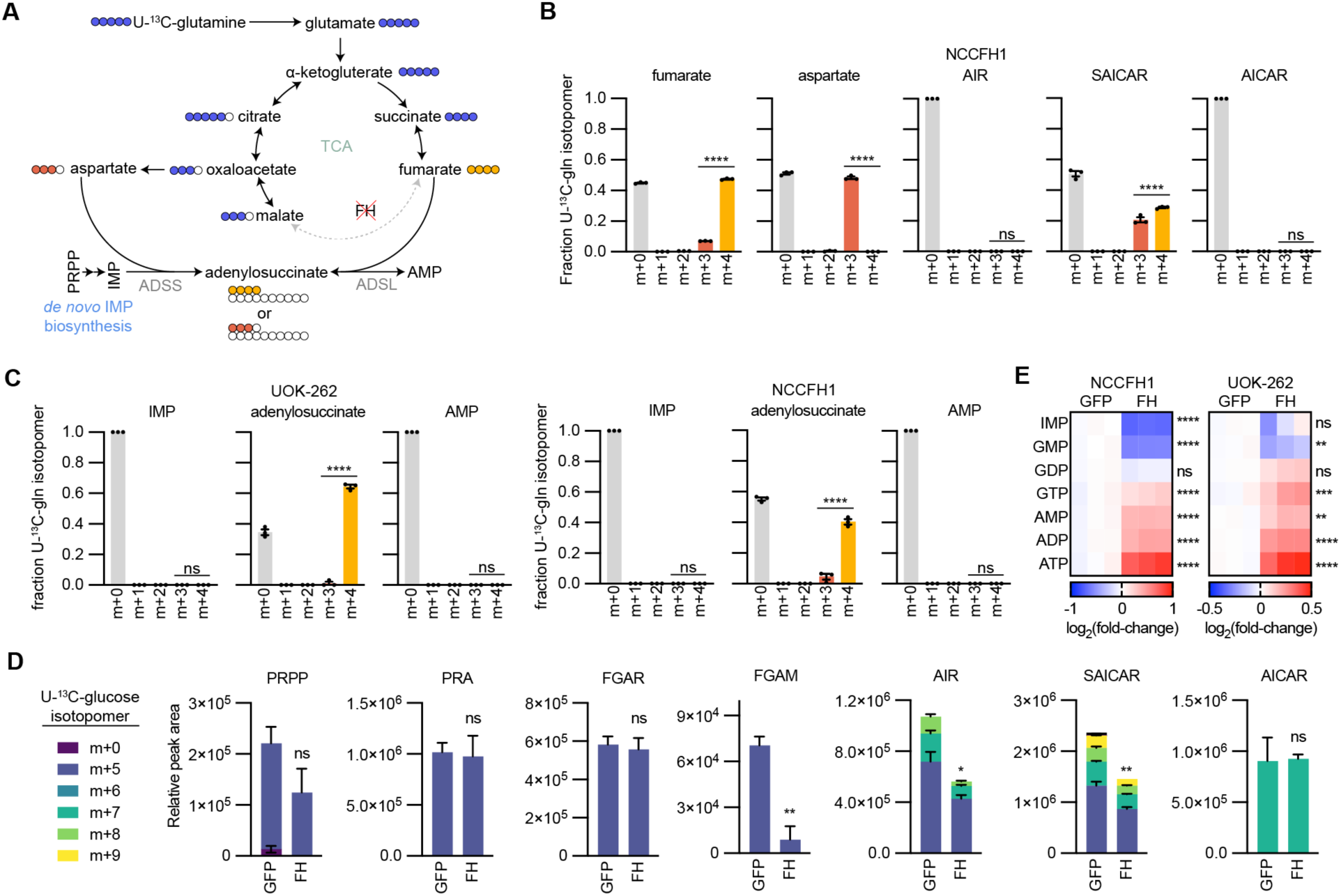
Loss of FH activity disrupts *de novo* purine biosynthesis. (**A**) Schematic representing the flow of U-^13^C-glutamine-derived carbons in the TCA cycle and AMP biosynthesis in HLRCC cells. Also depicted is purine synthesis via the salvage pathway. (**B**) Fraction of ^13^C-labeled isotopologues for fumarate, malate, AIR, SAICAR, and AICAR in UOK-262 cells were treated with U-^13^C-glutamine. (**C**), Fraction of ^13^C-labeled isotopologues for IMP, adenylosuccinate, and AMP in HLRCC cell lines. (**D**) Relative peak areas for *de novo* IMP biosynthesis intermediates in NCCFH1 cells treated with U-^13^C-glucose. Bars are colored by the contribution of individual isotopologues. (**E**) log_2_(fold-change) for purine nucleotides in HLRCC cell lines expressing GFP or FH.

**Supplementary Fig. S14:**
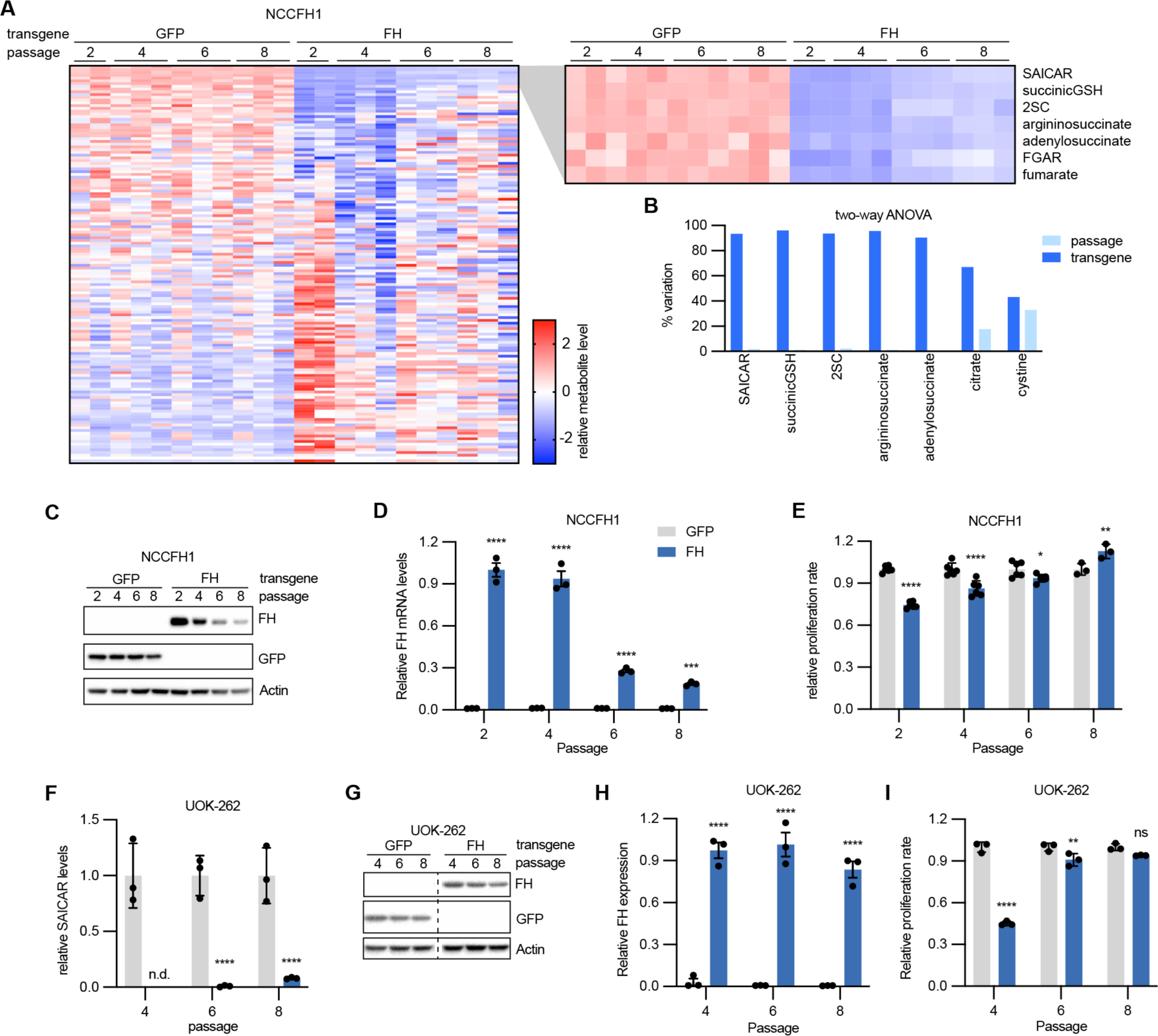
Dysregulated purine metabolism is independent of HLRCC cell proliferation rate. NCCFH1 cells were infected with lentivirus encoding GFP or FH and analyzed for 8 passages and analyzed for (**A**-**B**) 148 metabolites, (**C**) FH protein, (**D**) FH mRNA, and (**E**) proliferation rates. UOK-262 were infected with lentivirus encoding GFP or FH and analyzed for 8 passages and analyzed for (**F**) SAICAR, (**G**) FH protein, (**H**) FH mRNA, and (**I**) proliferation rates.

**Supplementary Fig. S15:**
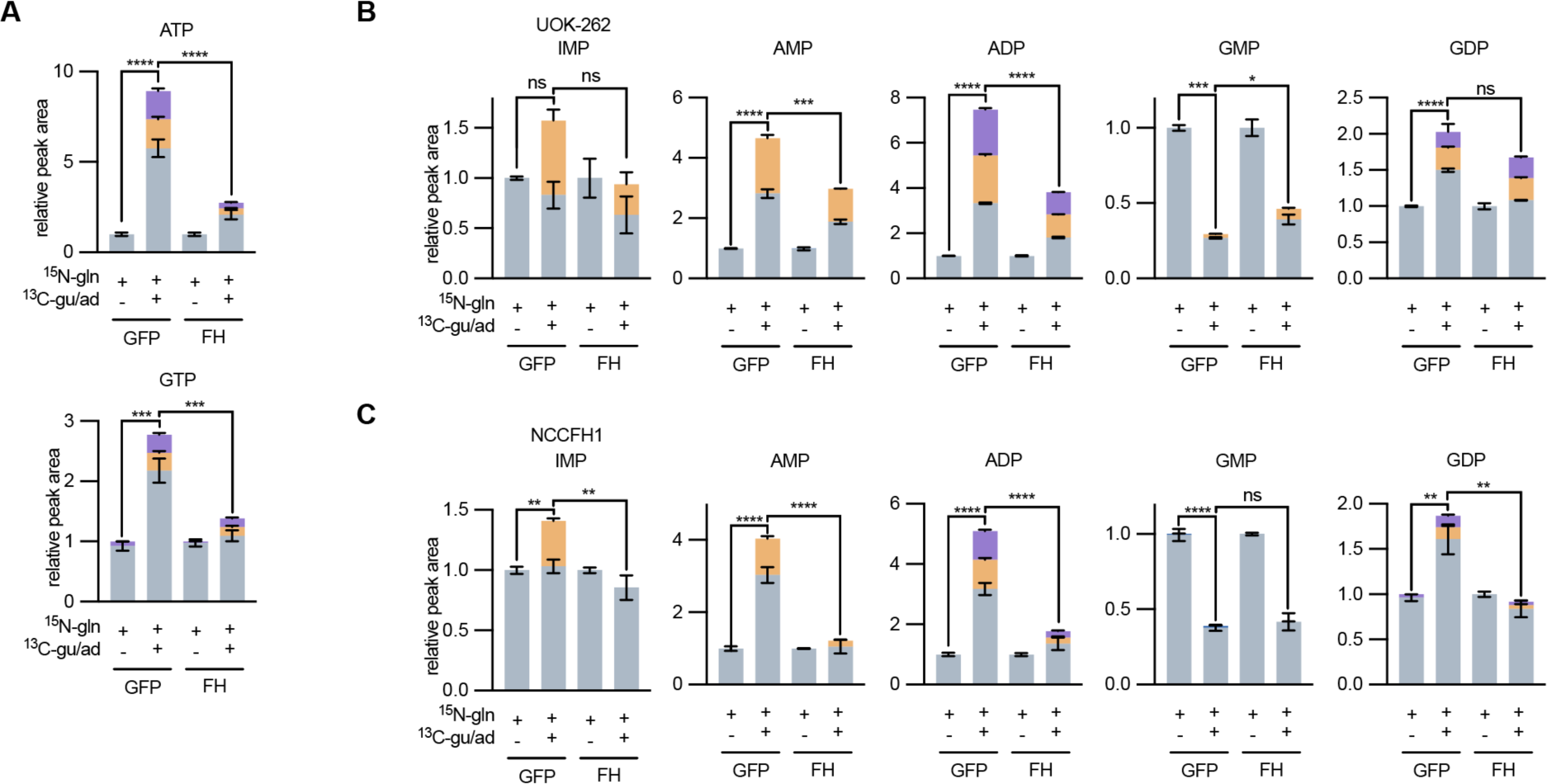
FH-deficient cells use the purine salvage pathway to maintain purine nucleotide levels. (**A**) NCCFH1 cells treated with for 6 hours with 4 mM amide-^15^N-glutamine, 50 µM 8-^13^C-adenine, and 50 µM 8-^13^C-guanine and ATP and GTP were analyzed by LC-MS for mass shifts corresponding to *de novo* synthesis (purple) or salvage pathway (orange). (**B**) UOK-262 cells treated with for 6 hours with 4 mM amide-^15^N-glutamine, 50 µM 8-^13^C-adenine, and 50 µM 8-^13^C-guanine and IMP, AMP, ADP, GMP, and GDP were analyzed by LC-MS for mass shifts corresponding to *de novo* synthesis (purple) or salvage pathway (orange). (**C**) NCCFH1 cells treated with for 6 hours with 4 mM amide-^15^N-glutamine, 50 µM 8-^13^C-adenine, and 50 µM 8-^13^C-guanine and IMP, AMP, ADP, GMP, and GDP were analyzed by LC-MS for mass shifts corresponding to *de novo* synthesis (purple) or salvage pathway (orange).

**Supplementary Fig. S16:**
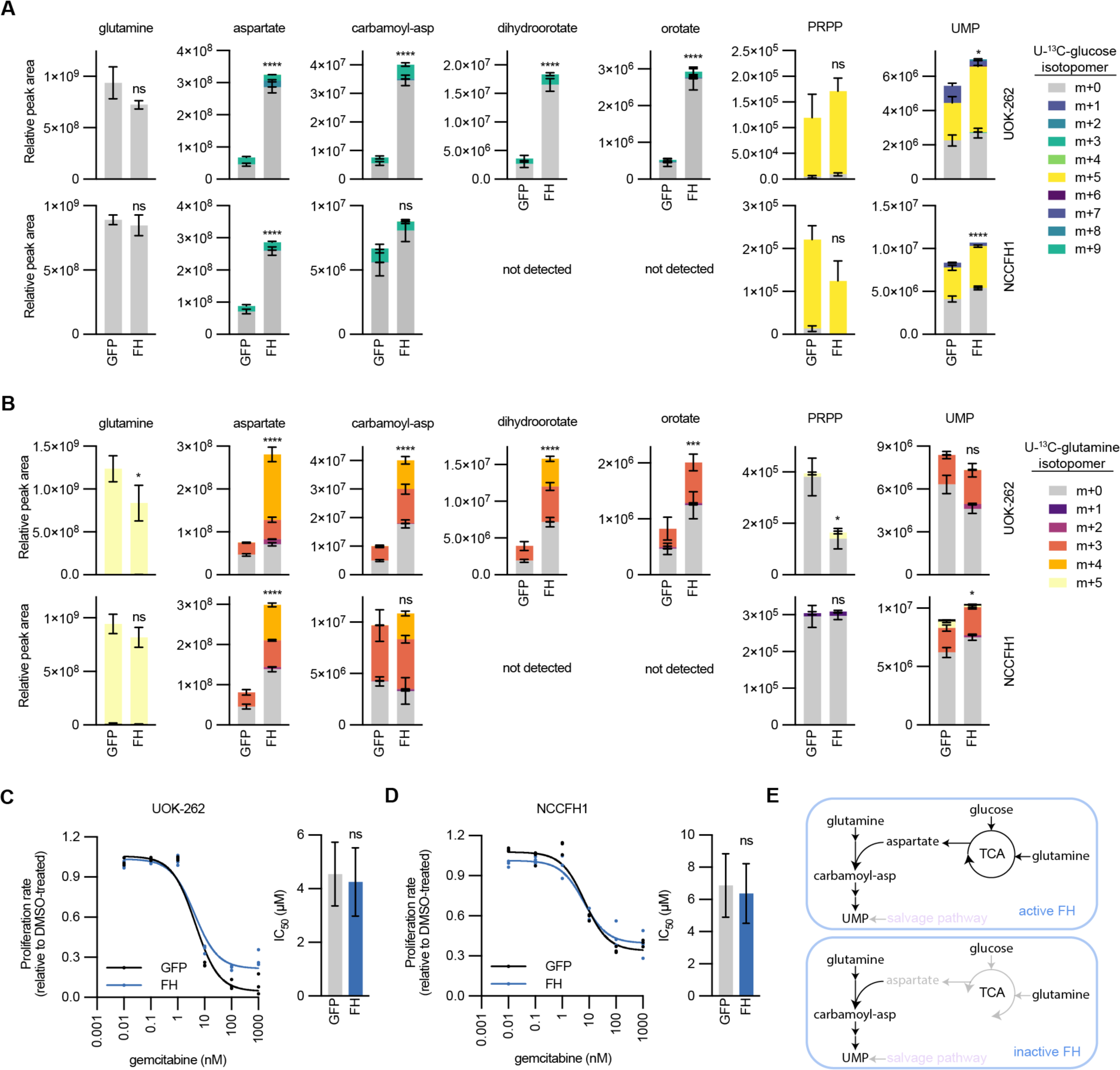
Pyrimidine metabolism in FH-deficient and FH-expressing cells. (**A**) Relative metabolite glutamine and pyrimidine biosynthesis intermediates in HLRCC cells expressing GFP or FH and treated with U-^13^C-glucose. Bars are colored by the relative abundance of isotopomers. (**B**) Relative metabolite glutamine and pyrimidine biosynthesis intermediates in HLRCC cells expressing GFP or FH and treated with U-^13^C-glutamine. Bars are colored by the relative abundance of isotopomers. (**C**-**D**) Proliferation (relative to DMSO treated cells) of NCCFH1 and UOK-262 cells expressing GFP or FH and treated with varying concentrations of gemcitabine. Fitted values for EC_50_ are higher for FH-expressing cells. (**E**) Schematic depicting the metabolism of pyrimidine synthesis intermediates in FH-deficient and FH-expressing cells.

**Supplementary Fig. S17:**
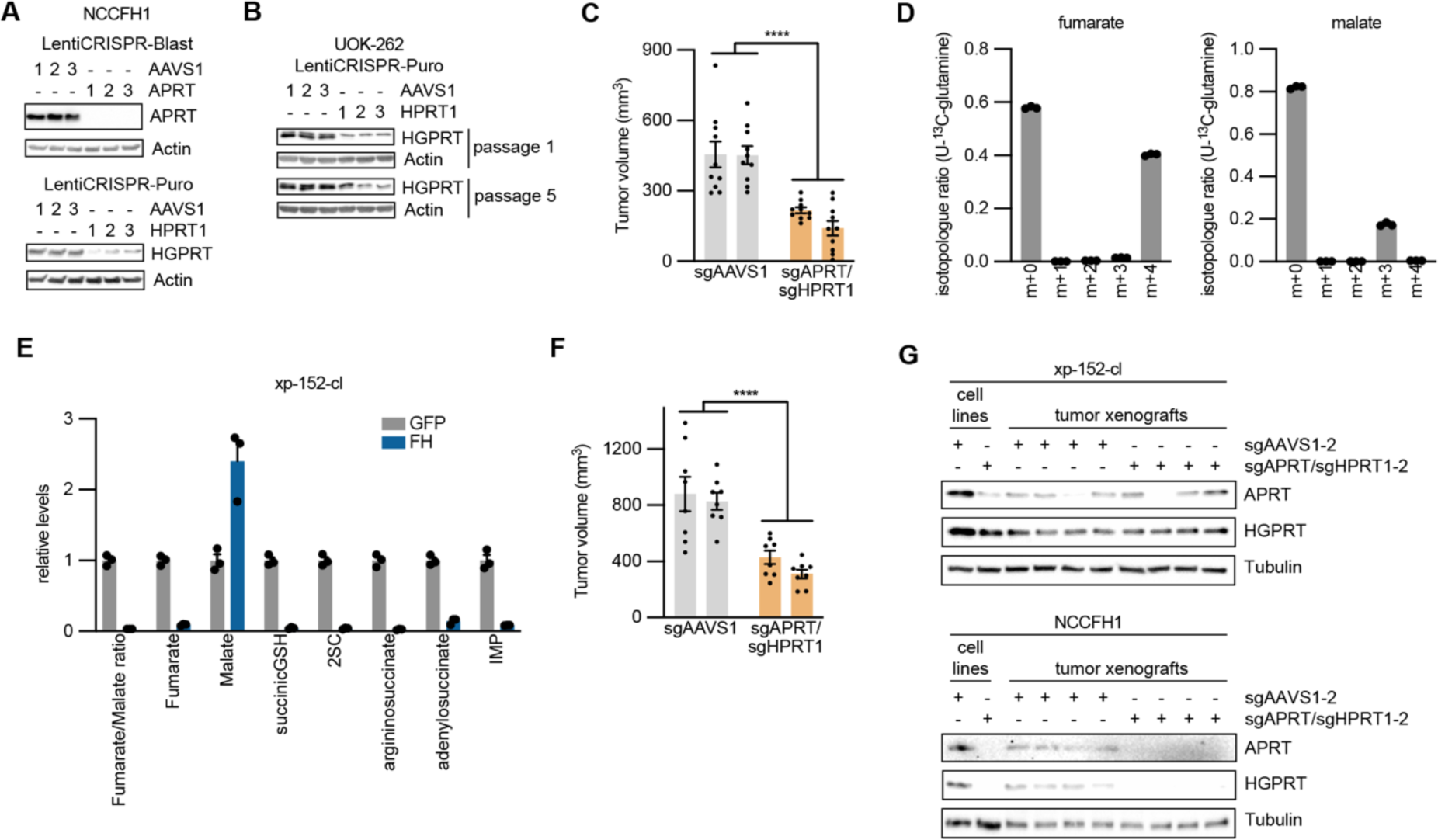
Disrupting expression of purine salvage enzymes impairs tumor growth. (**A**) Immunoblot analyzing APRT and HGPRT levels in NCCFH1 cells infected with LentiCRISPR-Blast with sgRNAs targeting AAVS1 or APRT (top) or infected with LentiCRISPR-Puro with sgRNAs targeting AAVS1 or HPRT1. (**B**) Immunoblot analyzing HGPRT levels in UOK-262 cells infected with LentiCRISPR-Puro with sgRNAs targeting AAVS1 or HPRT1 at passages 1 and 5 post-infection and selection. (**C**) Endpoint tumor volumes of NCCFH1 tumor xenografts expressing sgAAVS1 or sgAPRT/sgHPRT1. (**D**) Fraction fumarate and malate isotopologues in xp-152-cl cells treated with U-^13^C-glutamine for 6 hours. (**E**) Relative levels of fumarate, malate, fumarate/malate ratio, and fumarate-regulated metabolites in xp-152-cl cells expressing GFP or FH. Endpoint tumor volumes of NCCFH1 tumor xenografts expressing sgAAVS1 or sgAPRT/sgHPRT1. (**F**) Immunoblots evaluating the expression of APRT and HGPRT in lysates from NCCFH1 and xp-152-cl cell lines and corresponding tumor xenograft lysates.

**Supplementary Fig. S18:**
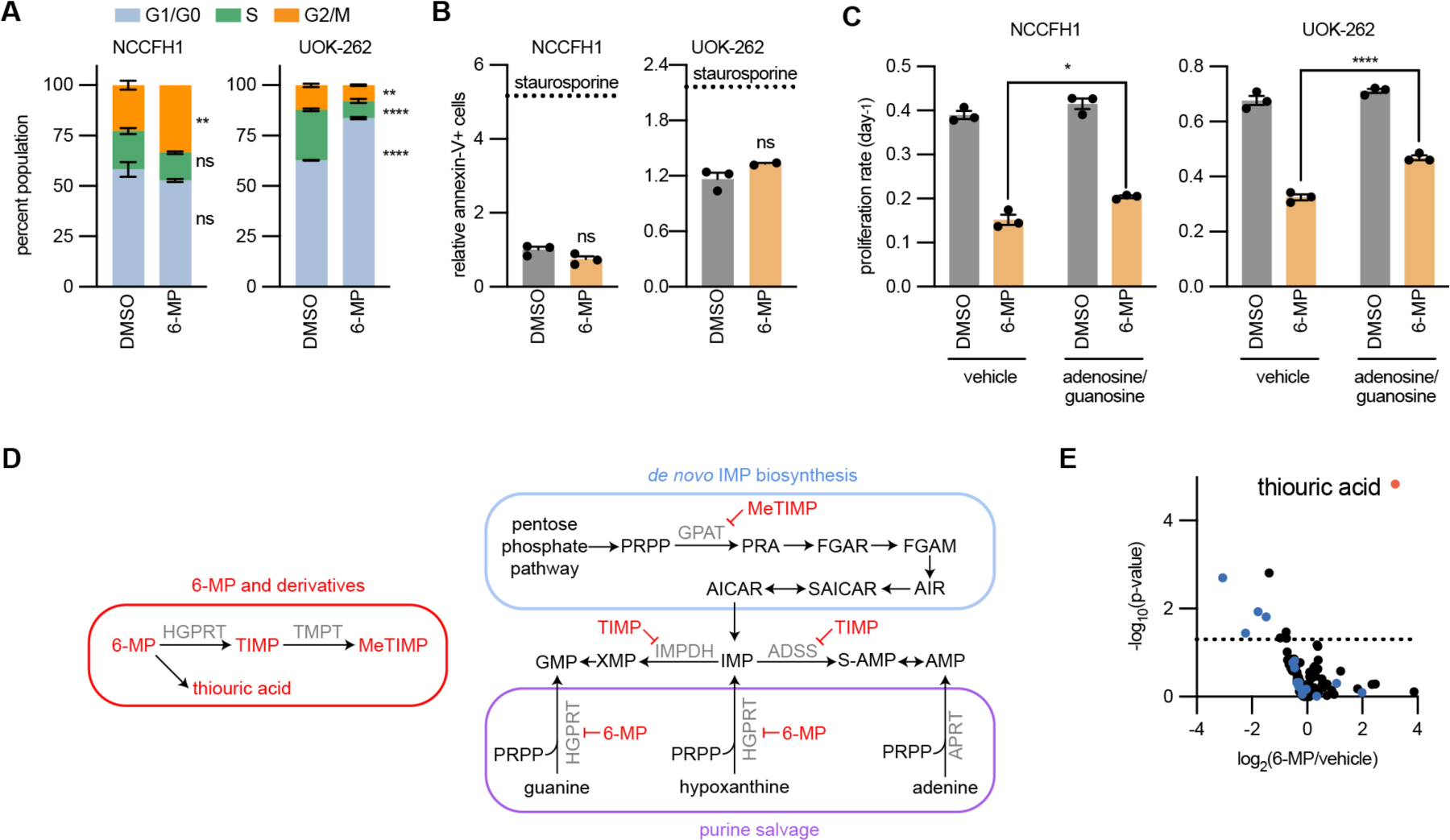
Purine salvage inhibitor 6-mercaptopurine (6-MP) impairs tumor growth. UOK-262 and NCCFH1 cells were treated for 48 hours with media containing DMSO or 100 µM 6-MP or for 24 hours with 0.1 µM staurosporine and analyzed for (**A**) proportion of cells within the cell cycle stages and (**B**) relative number of cells undergoing apoptosis. (**C**) Proliferation rates of UOK-262 and NCCFH1 cells treated with media containing 100 µM 6-MP, 10 µM adenosine, and 10 µM guanosine. (**D**) Schematic depicting how 6-MP disrupts purine salvage and biosynthesis. 6-MP is a competitive inhibitor of HGPRT. It is metabolized by HGPRT to generate 6-thioinosine-5-monophosphate (TIMP), which inhibits IMPDH and ADSS in GMP and AMP synthesis respectively. Further, TIMP can be metabolized by TMPT to generate MeTIMP, which inhibits GPAT, which catalyzes the initiating step in *de novo* IMP synthesis. (**E**) Volcano plot depicting the significant changes in NCCFH1 tumor xenografts from mice treated with 40 mg/kg 6-MP. We were able to detect thiouric acid, which accumulated in the tumors of 6-MP treated mice. Blue dots represent purine nucleotides and purine biosynthesis intermediates.

## Notes

Conflict of Interest: The authors declare no potential conflicts of interest.

### Competing Interest Statement

The authors have declared no competing interest.

### Summary of Updates

Addition of many pieces of in vitro and in vivo data supporting the conclusion that FH-deficient cells rely on purine salvage to maintain purine nucleotide levels and tumor growth.

